# Seasonal migration alters energetic trade-off optimization and shapes life history

**DOI:** 10.1101/2023.09.12.557441

**Authors:** Allison K. Pierce, Scott W. Yanco, Michael B. Wunder

**Affiliations:** University of Colorado Denver, Department of Integrative Biology, Denver, CO 80217, USA; Center for Biodiversity and Global Change, Yale University, New Haven, CT 06520, USA; Department of Ecology and Evolutionary Biology, Yale University, New Haven, CT 06520, USA

**Keywords:** migration, pace of life, life history, trade-offs, energy budget, movement, evolution, seasonality, DEB

## Abstract

Trade-offs between current and future reproduction manifest as a set of co-varying life history and metabolic traits, collectively referred to as “pace of life” (POL). Seasonal migration modulates realized environmental dynamics and putatively affects realized POL, however, the mechanisms by which migratory behavior shapes POL remain unclear. We explored how migratory behavior interacts with environmental and metabolic dynamics to shape POL. Using an individual based model of movement and metabolism we compared fitness-optimized trade-offs among migration strategies. We found annual experienced seasonality and migration distance primarily drove POL differentiation through developmental and migration phenology trade-offs. Similarly, our analysis of empirically-estimated metabolic data from 265 bird species suggested seasonal niche conservatism and migration distance interact to drive POL. We show multiple viable life history strategies are conducive to a migratory lifestyle. Overall, our findings suggest metabolism mediates complex interactions between behavior, environment, and life history.

**Authorship statement:** AKP and SWY contributed equally to this work. AKP and SWY conceived the project, designed the model, performed analyses, and drafted the manuscript with support from MBW. Simulation and optimization model code written by AKP with input from SWY and MBW. Empirical data analysis and code lead by SWY with input from AKP and MBW. All authors contributed critical input to the manuscript.

**Data accessibility statement:** All R code for models and analyses are available at https://doi.org/10.5281/zenodo.10145976 and https://github.com/allipierce/Pierce_et_al_2023_MigrationPOL. The empirically derived data used in this work are all publicly available. DEB data are available from the Add-My-Pet portal at https://www.bio.vu.nl/thb/deb/deblab/add_my_pet/. Geographic range data are available from BirdLife International at http://datazone.birdlife.org/species/requestdis. Niche dissimilarity data came from https://doi.org/10.1111/geb.13722 and phylogeny data are available from VertLife at http://vertlife.org/data/.

## Introduction

Time and energy are limited resources used by organisms to survive and reproduce (Levins 1962; Cody 1966). Organisms balance use of these resources by varying strategic emphasis on current versus future reproduction (Williams 1966; van Noordwijk & de Jong 1986; Stearns 1989, 1992). This fundamental trade-off manifests as a set of co-varying life history and metabolic trade-offs, collectively referred to as ‘pace of life’ (POL; Williams 1966; Stearns 1989, 1992; Wikelski & Ricklefs 2001; Ricklefs & Wikelski 2002) driven by traits which can include age at first reproduction (Harvey & Zammuto 1985; Sæther 1988), fecundity (Lack 1947; Sæther 1988), growth rate (Gebhardt-Henrich & Richner 1998), investment in survival (Harvey & Zammuto 1985), and metabolic rate (Wikelski *et al*. 2003a; Wiersma *et al*. 2007b).

At biogeographic scales, latitude covaries with both life history trade-offs (Moreau 1944; Lack 1947; Martin 1996; Ricklefs 2000; Jetz *et al*. 2008; Yanco *et al*. 2022) and organismal metabolism (Wikelski et al. 2003a; Wiersma et al. 2007a, b; Yanco et al. 2022) implying that environment contributes to realized POL. However, despite the extensive literature on the topic, the mechanistic drivers of covariation between environment and POL remain poorly understood, largely because latitude is actually a proxy for several confounded variables (e.g., mean resource availability, periodic amplitude of resource variance, ambient temperature, predation pressure, intraspecific competition, etc.) all of which could plausibly drive variance in POL (Ghalambor & Martin 2001). Yanco et al. (2022) showed that high amplitude seasonal variation in resource availability mechanistically constrains organisms to relatively “fast” POL strategies (emphasis on fecundity over survival, high metabolic rate). Conversely, low amplitude seasonal resource fluctuation was associated with expanded viability of “slow” POL strategies (emphasis on survival over fecundity, low metabolic rates). These findings imply that environmental conditions, and the dynamics of those conditions over time, influence realized POL strategies.

POL also exhibits complicated covariance patterns with animal behavior. Behavior can modify extrinsic mortality risk, which is expected to drive POL; High extrinsic mortality risk favors a fast POL whereas low extrinsic mortality risk favors a slow POL (Ricklefs 2000; Réale *et al*. 2010; Dammhahn *et al*. 2018). Behaviors such as movement may also mediate the environmental conditions an organism experiences, making the optimal life history strategy the outcome of an interaction between environment and behavior rather than environment alone (Spiegel *et al*. 2015; Campos-Candela *et al*. 2019; Laskowski *et al*. 2021). For example, populations of Corsican Blue Tits (Cyanistes caeruleus) in evergreen-dominated habitats exhibited slow POL along with reduced handling aggression, exploration speed, and nest defense whereas individuals in deciduous habitats showed the converse (Dubuc-Messier *et al*. 2017).

Seasonal migration is a behavior with putatively substantial effects on realized POL but empirical findings about the direction and magnitude of those effects vary depending on the taxonomic scale of comparison, environmental context, and characterization of the behavior (Soriano-Redondo *et al*. 2020; Winger & Pegan 2021). Migration radically alters the environmental conditions an organism experiences, typically leading to niche conservatism, relatively invariant conditions across the annual cycle (Nakazawa *et al*. 2004; Gómez *et al*. 2016; Merkle *et al*. 2016; Thorup *et al*. 2017; Zurell *et al*. 2018; Alerstam *et al*. 2019; Abrahms *et al*. 2021; Yanco *et al*. 2021), which might be expected to favor a slow POL. On the other hand, migration is a potentially costly endeavor both in terms of increased mortality risk (Sillett & Holmes 2002; Rushing *et al*. 2016, 2017; but see Zúñiga *et al*. 2017) and metabolic expenditures (Wikelski *et al*. 2003b; Alves *et al*. 2013; Brown *et al*. 2023), suggesting that migration might be associated with a fast POL. Intriguingly, some studies suggest that migrant birds exhibit intermediate POL: faster than temperate counterparts, but slower than residents breeding at similar latitudes. For example, avian clutch size is a classic case study of biogeographic variance in life history across a gradient of seasonality (Moreau 1944; Lack 1947). Jetz et al. (2008) showed that clutch sizes of migrants were smaller than would be expected from phylogenetically equivalent residents breeding at similar latitudes. Similarly, Weirsma et al. (2007b) showed that although tissue basal metabolic rate (BMR; assumed to be proportional to POL) was positively related with latitude, migrants had lower BMR than their resident temperate counterparts and higher BMR than tropical residents. Ultimately, how migration is mechanistically linked to realized POL remains ambiguous because, to date, few studies have simultaneously accounted for the complex feedbacks between the bioenergetics of migration and the emergent environmental dynamics.

In this study we sought to explore how migratory behavioral mitigation of seasonality can affect dynamics of metabolic and temporal trade-offs to shape POL and life history based on a model of avian migration. For example, if migration solely balances the energetic cost of movement against the benefit from mitigating seasonal resource variance, we would expect similar life histories across migratory strategies. However, if migration substantially affects the net energy available to balance metabolic tradeoffs, then we would expect covariation between migratory strategy and life history. Furthermore, energetic trade-offs also interact with phenological constraints imposed by migration. Time-consuming migrations might restrict breeding season duration and ultimately favor fewer and/or faster developing young despite increases in net energy from migration. Thus, certain migrations may be more time- than energy-limited than others.

To model energetic and temporal trade-offs we combined an individual-based model (IBM) of one-dimensional movement within a simulated environment of seasonally fluctuating energy with a generalized Dynamic Energy Budget (Kooijman 2010) model of metabolism (sensu Yanco *et al*. 2022). We compared life-history optimization trade-offs between migratory and stationary strategies using a heuristic optimization algorithm to solve for fitness-optimized IBM parameterizations across a gradient of seasonality and migration strategies. Finally, we show empirically that seasonal niche conservatism and migration distance interact to drive realized POL across 265 species of birds. We suggest that behavioral phenotype may play an underappreciated role in driving life history strategies, especially in cases where the interaction of behavior and the environment produce novel adaptive landscapes.

## Methods

### DEB model description

Dynamic Energy Budget (DEB) theory (Kooijman 2010) proposes a dynamic mechanistic model of how individuals uptake, use, and dissipate energy over their lifespans and across different life stages based on the principles of energy and mass conservation, thermodynamics, and stoichiometric constraints. To simulate energy acquisition and allocation dynamics of metabolic trade-offs we implemented a unitless specification of a standard DEB model modified to incorporate metabolic costs of migratory movements (Kooijman 2010). We scaled DEB parameters for energy and organism structural size by maximum reserve energy density and maximum length respectively. Resultant system dynamics are therefore more generalizable as they quantify proportional changes rather than absolute changes in units of energy or body size.

In the standard model (Fig. 1), organisms acquire energy from the environment and assimilate it into reserves (*ṗ_A_*). Energy is then mobilized from the reserve at a fixed proportion *κ* to cover energetic requirements of growth (*ṗ_G_*) and somatic maintenance (*ṗ_S_*), with the remainder diverted to maturation maintenance (*ṗ_J_*) and reproduction (*ṗ_R_*). In the juvenile stage, energy allocated to reproduction is applied to advancing maturity level rather than offspring production.

**Figure 1.**
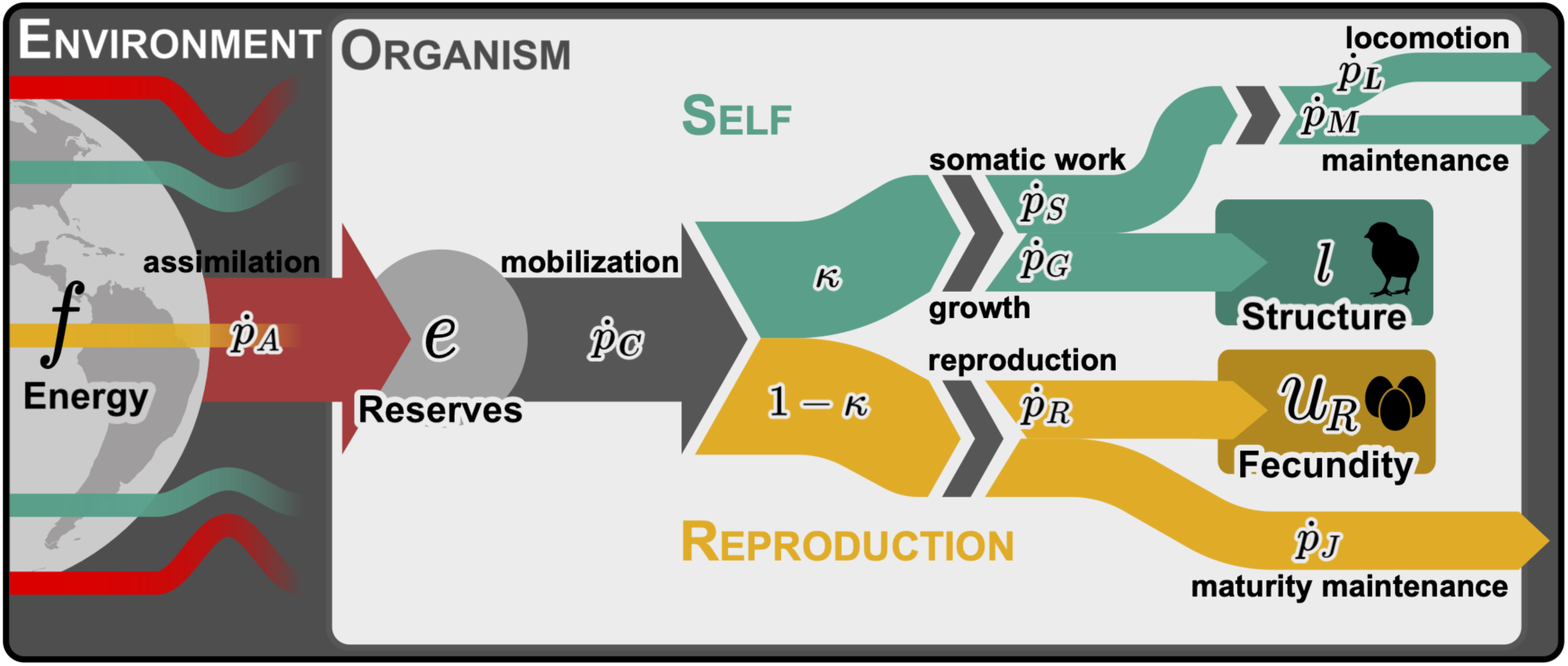
Dynamic energetic budget model flow diagram illustrating energy uptake and dissipation during the adult age stage where energy from the environment (*f*) is assimilated into the organism’s reserves (*e*) then mobilized for allocation to somatic and reproductive processes.

Locomotory costs are not explicitly defined but are assumed to come from somatic maintenance costs when they are independent from foraging rate (Kooijman 2010). Therefore, we modified the standard model to isolate migration movement costs from somatic maintenance costs via the compound parameter *m* which we defined as the ratio between metabolic cost of locomotion to travel one body length (*l*) during migration relative to somatic maintenance costs unrelated to migration. Thus, total somatic maintenance rate (*ṗ_S_*) is the sum of the locomotory cost of migration (*ṗ_L_*) which scales with structural surface area (*l^2^*) and distance, and somatic maintenance costs unrelated to migration (*ṗ_M_*) which are proportional to biovolume (*l^3^*).

In starvation conditions, energy use is prioritized by somatic maintenance first, growth second, followed by maturation/reproduction. In our implementation, length or maturation level cannot shrink to reduce energy costs and death occurs when energy drawn for somatic maintenance exceeds available reserves. Detailed model parameterization and derivation is available in supplement Appendix S1 and Kooijman (2010).

### IBM simulation

We used an IBM simulation to evaluate daily energy acquisition and allocation for an individual over 365 days from birth, through sexual maturation, to reproduction. At each time-step a set of differential equations was numerically solved to update values of the DEB state variables given individual trait parameters (Table 1; see Appendix S1 for detail). Environmental energy available at each time-step depended on the individual’s latitudinal location, where available energy *f* varied across 365 discrete time-steps as one oscillation of a sine function, for which the amplitude and phase varied spatially across the 180 continuous latitude units.

**Table 1.**
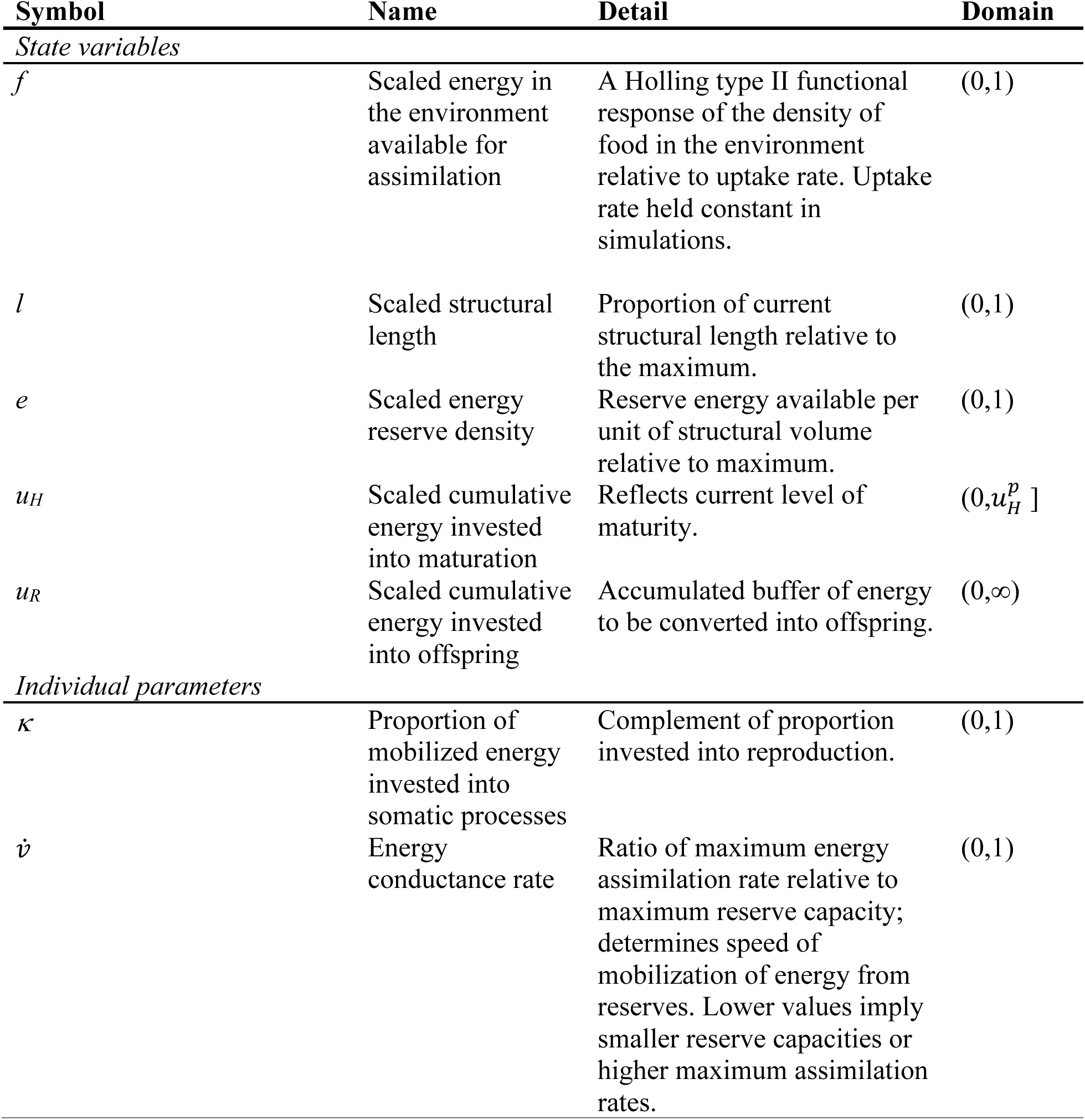
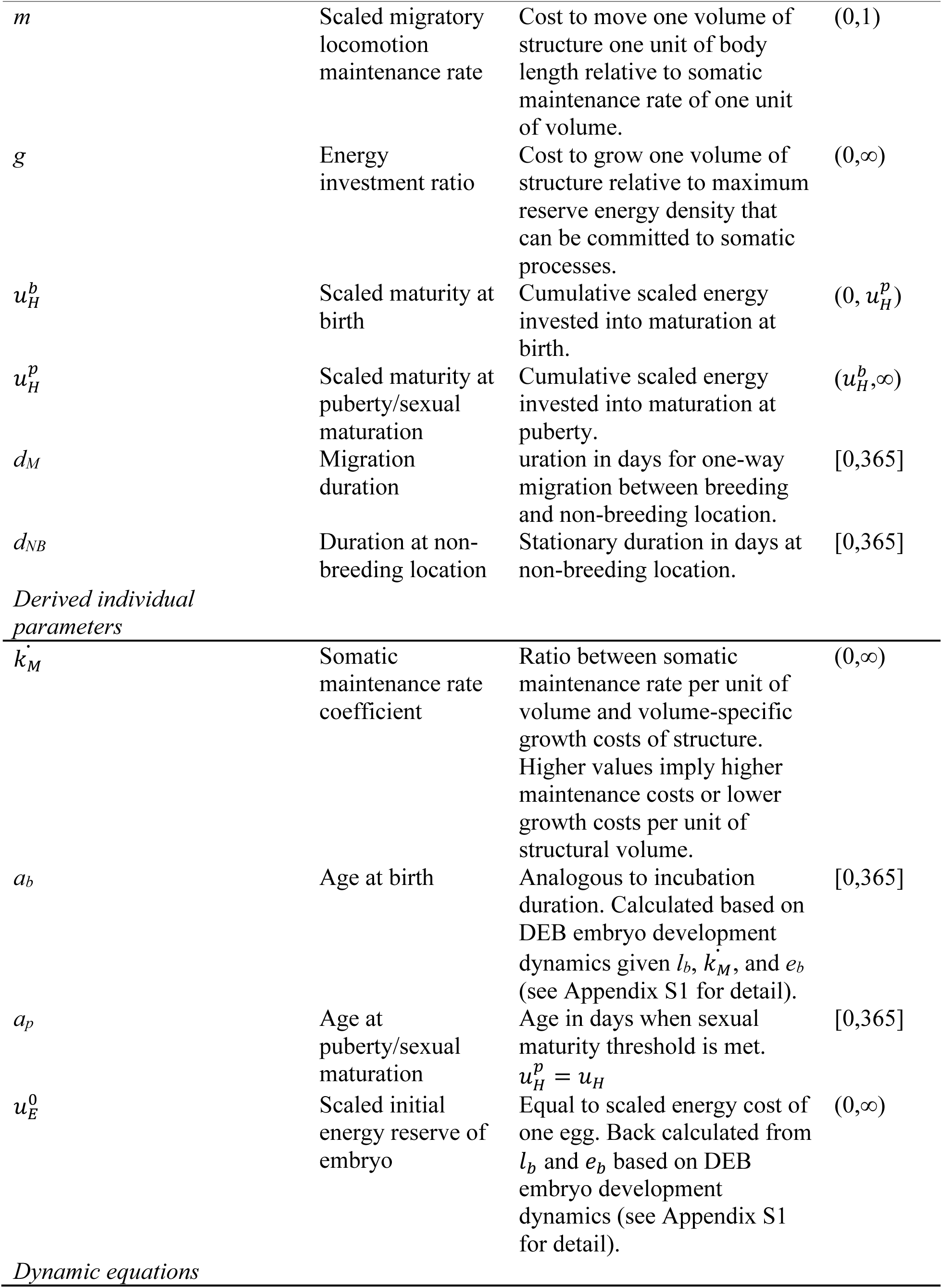
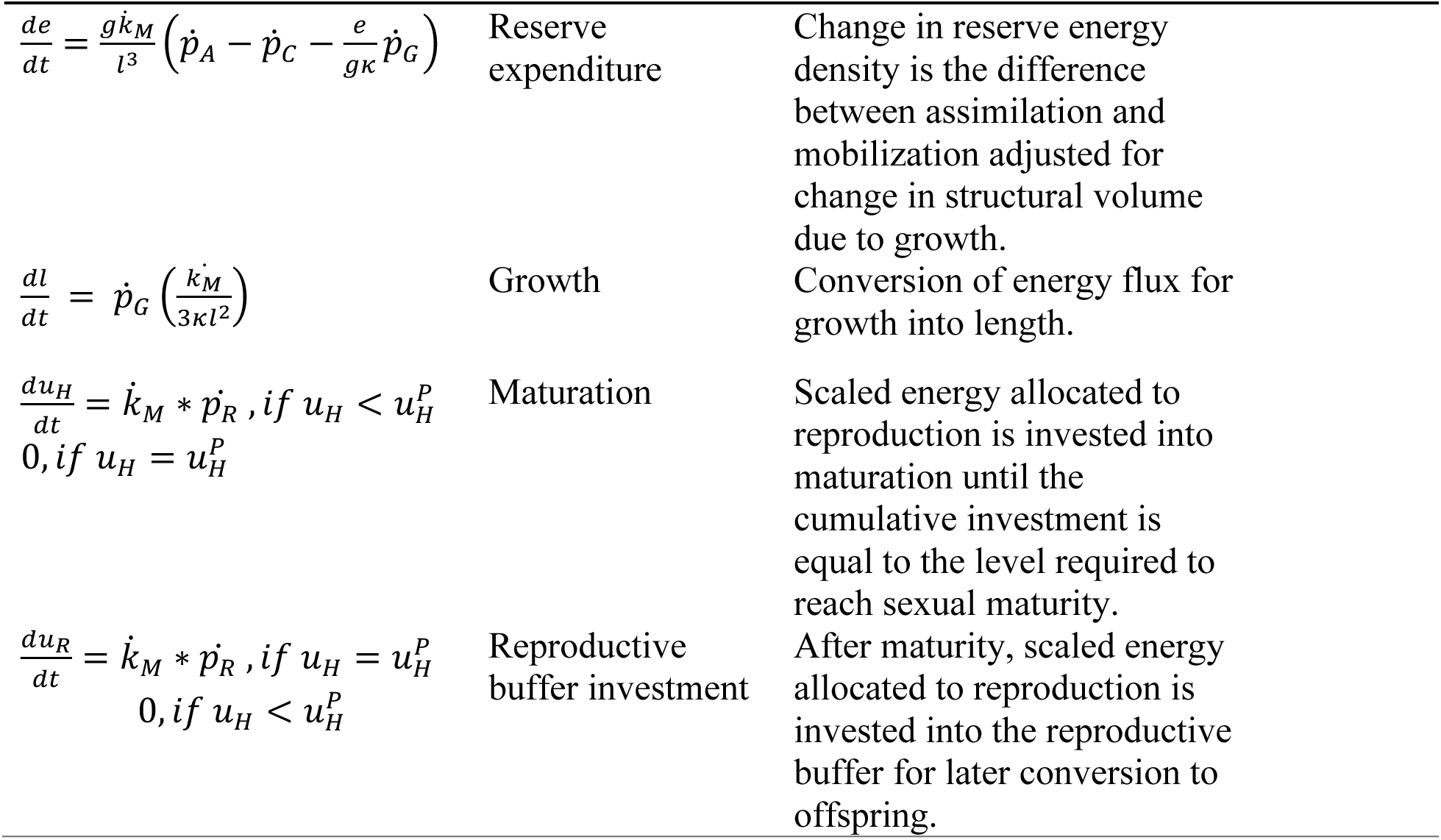
IBM state variables, individual parameters, and equations used to evaluate DEB and movement model dynamics with potential value ranges for each parameter. During the model optimization process we further restricted value ranges for some parameters to account for covariation between parameters and to reduce generation of inviable solutions (see supplement Appendix S1 for detail).

Maximum amplitude occurred at the lowest and highest latitudes and decreased towards the mid-point. Phase is shifted by a half wavelength on either side of the mid-point to reflect the alternating seasonality between hemispheres (sensu Yanco et al. 2022).

Individuals are initiated at the peak of seasonal energy availability for their initial location and reproduction occurs such that offspring are born when energy availability peaks again on day 365 (*t* = 365 - *a_b_*). If individuals are migratory, at the end of each time-step they move towards non-breeding locations once sexually mature or depart to breeding locations after remaining at their non-breeding location for their specified duration (*d_NB_*). For simplicity, we assumed constant migration speed. Distance traveled in each time-step was calculated as migration distance divided by specified duration of migration (*d_M_*).

We calculated fecundity for viable solutions as the amount of cumulative energy in the reproductive buffer divided by the cost of one egg, *u*_E_^0^. Egg cost is based on the individual’s scaled reserve density at egg production with the assumption that length at birth and growth rate of offspring are inherited traits.

At the final time-step, solutions (individual sets of parameters) were classified as nonviable if energy reserves were depleted (*e* < 0), reserves at initialization were insufficient for offspring to reach birth size (*e_b_* < *l_b_*), less than one offspring was produced, or migratory individuals did not return to breeding locations in time to complete reproduction.

### IBM parameter optimization

Heuristic optimization algorithms are useful for efficiently solving multi-dimensional optimization problems that are analytically intractable and too computationally intensive to solve numerically by brute force (Hamblin 2013), such as the problem we studied here. We maximized for fecundity using a genetic algorithm, a heuristic optimization of an objective function (represented here as an IBM) using iterative search techniques inspired by genetic mechanisms of natural selection such as inheritance, mutation, and crossover. Despite the biological inspiration for the algorithm, our implementation is not intended to be interpreted as simulating evolutionary processes.

### Optimization by movement strategy

To investigate life history optimization across a gradient of seasonality and migration distance we optimized parameterizations for 9 different movement strategies: 6 migratory and 3 stationary. Stationary strategies occurred at latitudes we classified as high seasonality (80, -80), mid seasonality (40, -40) and aseasonal (0). Migratory strategies were defined by breeding and non-breeding end point locations and varied in distance such that migrants bred in high or mid-seasonal latitudes and migrated to high or mid-seasonal locations in the opposite “hemisphere”, a mid-seasonal location in the same hemisphere, or the aseasonal location.

Migration strategies were further classified by migration distance, seasonality of endpoints, and energy management strategy. Migrations between seasonally out of phase endpoints (cross hemisphere) were defined as energy-buffering strategies because they buffer seasonal oscillations of energy over time by tracking seasonal peaks in energy availability. We classified migrations between seasonally in-phase locations (within hemisphere) as energy mitigating strategies because they dampen amplitude of seasonal oscillations in energy over time. Stationary strategies do not behaviorally manipulate seasonality and so we classified them as coping strategies.

For each strategy, we initialized IBM parameters with randomly generated values for 5000 solutions and iteratively optimized values for 200 iterations (see Appendix S1 for detail). We repeated this process 1000 times (runs) to ensure adequate exploration of the solution space. Parameters were not simultaneously optimized across migratory strategies, so different strategies might have explored the solution space to different extents (Hamblin 2013). To address this, we used the same random number generation seeds across migration strategies for each run so that each optimization process started with the same set of strategy invariant parameters for each strategy but with different random sets for each run. We also performed an additional set of 90 runs per migratory strategy pre-initialized with parameters from five of the highest fecundity viable solutions for each strategy from the initial 1000 runs (10 runs of 9 pre-initializations) to equalize exploration of the solution space between strategies. Finally, we pooled together solutions from all 1090 runs, removed duplicate solutions within strategies, and retained solutions in the top 1% for fecundity from each for analysis.

We conducted all simulations, optimizations, and analyses in R (R Core Team 2022).

### Pace of Life Trade-offs

We used a correlation-based principal components analysis (PCA) of optimized individual parameter values from each movement strategy to analyze trait covariation patterns and evaluate metabolic and temporal trade-off dynamics. We analyzed energy mobilization and somatic allocation parameters *v̇* and *κ*, in addition to parameters governing downstream somatic and reproductive allocation (*k̇_M_*, *m*, and *g*). Additionally, we analyzed parameters related to timing of migration (*a_p_*, *d_NB_*, and *d_M_*). We used bi-plots of parameter loadings to qualitatively evaluate covariance relationships (especially including trade-offs) between parameter combinations and between migration strategies.

### In silico experiment

To investigate the effects of migration on relative life-history optimization across migratory strategies, we evaluated the fecundity values of optimized parameterizations for each migration strategy in simulation scenarios wherein individuals were forced to stay in breeding locations (could not migrate) as compared to optimized parameterizations for stationary strategies from the same respective breeding location.

### Empirical Analysis

We leveraged publicly available datasets to explore the extent of empirical support for the influence of migratory phenotype, biogeography, and realized niche dynamics on life history in birds. We used the Add-my-Pet dataset which includes empirically estimated DEB parameters for 4035 species (Marques *et al*. 2018) to calculate a general proxy for POL as a size-normalized estimate of relative maximum somatic mobilization rate *κṗ_C_* where higher values represent increased investment of reserves into somatic processes and thus slower pace of life. To account for non-linear scaling of energy assimilation and mobilization as a function of body size, we calculated this as 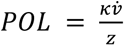, where z is the ratio of species-specific maximum length to the maximum length of 1 cm for a theoretical reference species, allowing for comparisons across species. We used this POL metric as the response variable in the models described below.

To assess the effects of biogeography, migratory phenotype, and environmental dynamics on these life history traits we considered three predictors: breeding latitude, migration distance, and environmental niche dissimilarity (roughly analogous to energy management strategy). We used the Birdlife International dataset (BirdLife International and Handbook of the Birds of the World 2016) to calculate the latitude of the centroid of breeding season ranges and calculated species-level migration distance as the great-circle distance between seasonal range centroids. To characterize seasonal environmental dynamics for each species we used results from Cohen & Jetz (2023) which estimates niche dissimilarity as the Mahalanobis distance in environmental space between niche hypervolumes of breeding locations and overwintering locations.

Combining these sources resulted in a final dataset of 265 species. All predictor variables were centered and scaled before model fitting.

We modeled macroecological patterns in our POL metric as a function of the three predictor variables under a Bayesian framework using the ‘brms’ package in R (Bürkner 2017). We modeled POL as beta distributed (expected domain [0,1]) with niche dissimilarity as a predictor on the beta distribution precision parameter (ϕ) based on previous reports (Yanco et al. 2022). To control for phylogenetic relatedness among the species in our sample we implemented a phylogenetic generalized mixed model approach (Lynch 1991; Garamszegi 2014). Briefly, this approach includes a random intercept by species and enforces a correlation structure among species matching a known phylogeny. We downloaded 100 phylogenies for our target species from vertlife.org (Jetz et al. 2012) and ran analyses using a single randomly selected tree. We re-ran analyses with new randomly selected trees to assess sensitivity to the specific phylogeny used and found results to be invariant across candidate trees. We fit the model using four MCMC chains of 10,000 draws (with a burn in of 5,000) thinned by 5. We assessed model convergence with a combination of *R̂* < 1.05 and visual inspection of MCMC traceplots and posterior predictive plots. We interpreted the relative significance of individual parameters using the Bayesian probability of direction (PD), the proportion of the estimated posterior distribution on the same side of zero as the estimate.

## Results

### Migration IBM

Movement paths of model optimized parameterizations for each migratory strategy generally tracked maximal available energy over time but all strategies optimized for later departure times from non-breeding locations than expected if perfectly tracking peak resources (Fig 2A). Migratory strategies realized higher annual mean energy availability than stationary strategies irrespective of breeding location with annual mean gain increasing with migratory distance (Fig. 2B).

**Figure 2.**
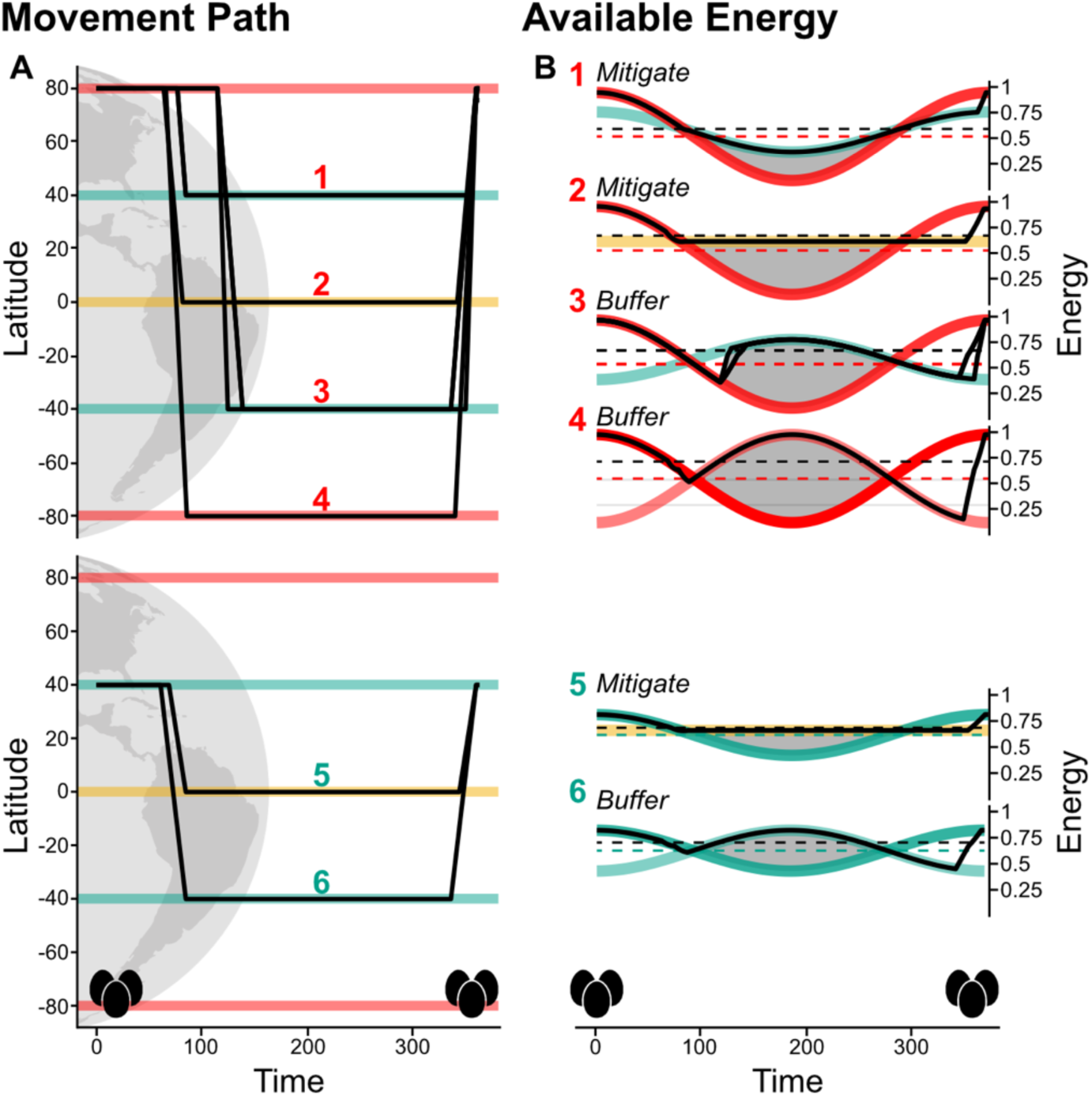
Model optimized migration movement paths (A) and available enroute energy (B) for migration strategies from breeding locations with high seasonality (1-4) and mid-seasonality (5,6) arranged by increasing migration distance top to bottom per location. In both panels, high seasonality is represented in red, mid in green, and aseasonality in yellow and egg images depict timing of birth and reproduction events. In panel A, solid black lines depict model optimized movement paths between latitudes of migration endpoints colored by seasonality for each strategy. In panel B, model optimized energy available enroute over time (black lines) overlay energy available in breeding locations (thick opaque red and green lines) and non-breeding areas (thick transparent lines colored by seasonality). Dashed lines depict mean annual energy available in breeding locations (red and green) and realized enroute (black). Shaded areas represent overwintering differences in energy availability between breeding and non-breeding locations.

### Pace of Life Trade-offs

Three axes of variation explained a total of 95.8% of the variation in optimized temporal and energetic trade-off parameters across all migratory and stationary strategies with components 1, 2, and 3 each explaining 58.3%, 27.4%, and 10.1% respectively. Secondary and tertiary somatic stream allocation parameters *k̇_M_*, *g*, and *m* were the primary contributors to the first component in addition to energy mobilization (*v̇*) and migration timing parameter *d_NB_* (PC1, Fig. S3). Somatic maintenance rate *k̇_M_* exhibited opposing loadings to all other parameters (*k̇_M_* = 0.44, *g* =-0.38, *m* =-0.43, *v̇* = -0.38, and *d_NB_*= -0.41) indicating trade-offs between somatic maintenance in metabolic cost of growth and migration as well as migration timing. Higher values of PC1 were associated with slower mobilization of reserves, prioritized investment into somatic maintenance over growth, and less time spent in the non-breeding location. Upstream energy allocation κ and parameters governing migratory timing *a_p_* and *d_M_* were the primary contributors of the second component (PC2, Fig. S3), all of which exhibited positive loadings (*κ*=0.58, *a_p_*=0.56, and *d_M_* =0.37) indicating higher values of PC2 were associated with higher proportional allocation to somatic processes, later maturation (and thus migration), and longer migration. Component 3 (PC3) was primarily associated with metabolic parameters *κ*, *g,* and *v̇* in addition to *a_p_* (Fig. S3). Loadings for *g* and *a_p_* (*g* =-0.83, *a_p_* =-0.38) opposed both *κ* and *v̇* (*κ*=0.41, *v̇*=0.42) indicating higher values of PC3 represent increased somatic allocation and mobilization of energy, lower investment into growth, and earlier maturation (and thus migration).

Optimized parameterizations were differentiated by energy management strategy along the PC1 and PC2 axes (Fig. 3A) indicating that trade-offs between metabolism and migratory timing were associated with behavioral manipulation of experienced seasonality (Fig. 3B).

**Figure 3.**
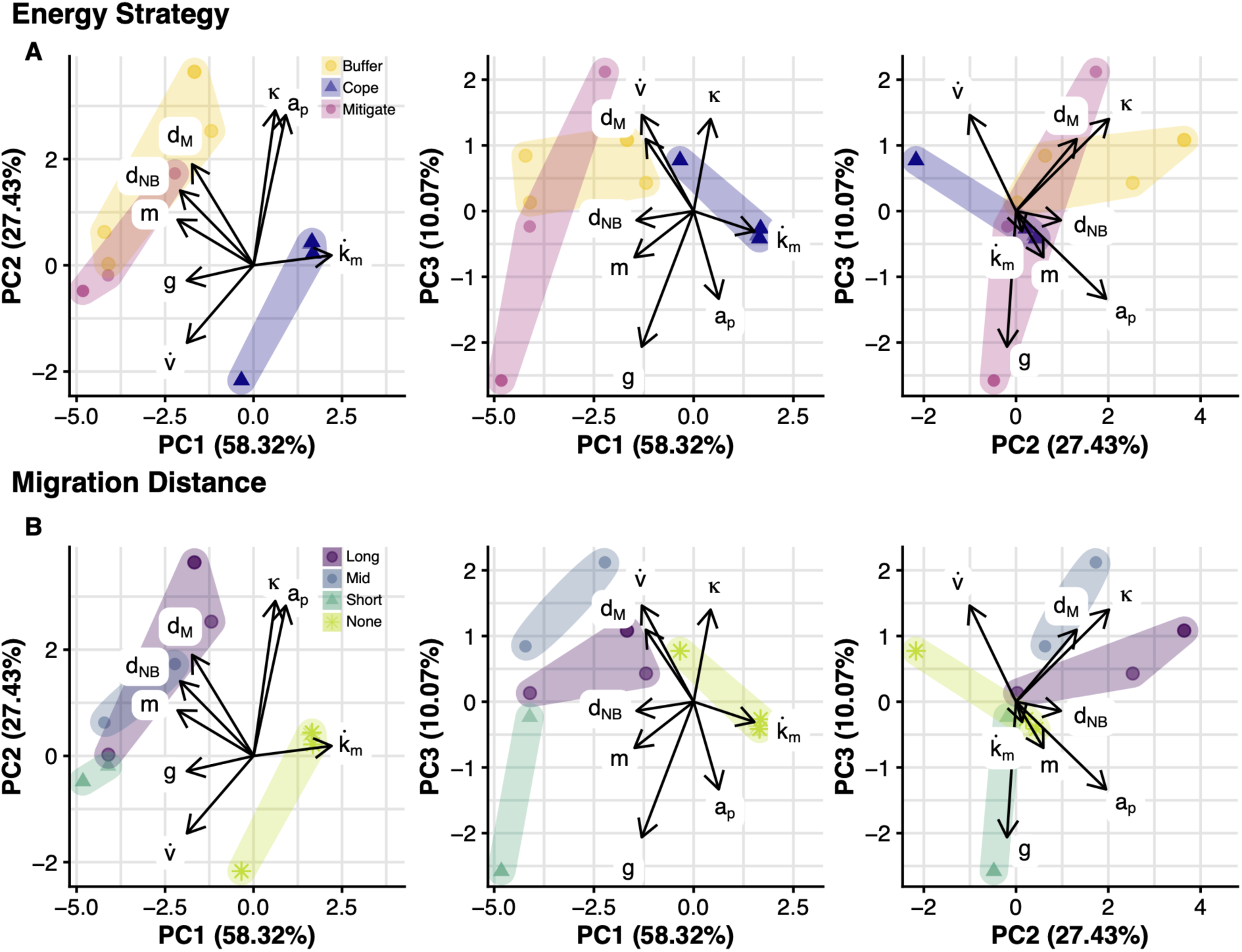
PCA of model optimized parameters depicting energetic trade-offs for stationary and migratory strategies differentiated by energy management strategy (A) and migration distance (B). Arrow length and direction indicate component loadings (correlations) of each parameter onto PCA axes.

Unsurprisingly and by design, stationary strategies that cope with seasonality differentiated from migratory strategies based on parameters for metabolic cost and timing of migration. More interestingly, stationary coping strategies were associated with lower per-volume growth costs, prioritized investment into somatic maintenance over growth, and slower mobilization. Buffering strategies were generally associated with longer migrations, shorter non-breeding stationary periods, and decreased migratory locomotion costs compared to mitigating strategies.

Migratory strategies were further differentiated by migration distance along the PC2 and PC3 axes (Fig 3B) such that somatic allocation and migration duration generally increased with distance and investment into growth and mobilization rate decreased. We did not observe appreciable differentiation with seasonality of breeding or non-breeding locations (Fig. S5).

### In silico experiment

Overall, energy-buffering long-distance migratory strategies realized the highest fecundity followed by the stationary aseasonal strategy. All optimized parameterizations for migratory movement strategies realized higher or similar relative fecundity than stationary strategies for their respective breeding locations with fecundity gain increasing with migratory distance. When model optimized parameterizations for migratory strategies were simulated to remain stationary at breeding locations, all parameterizations realized lower relative fecundity than optimized stationary parameterizations. Declines in relative fecundity generally increased with migration distance (Fig. 4).

**Figure 4.**
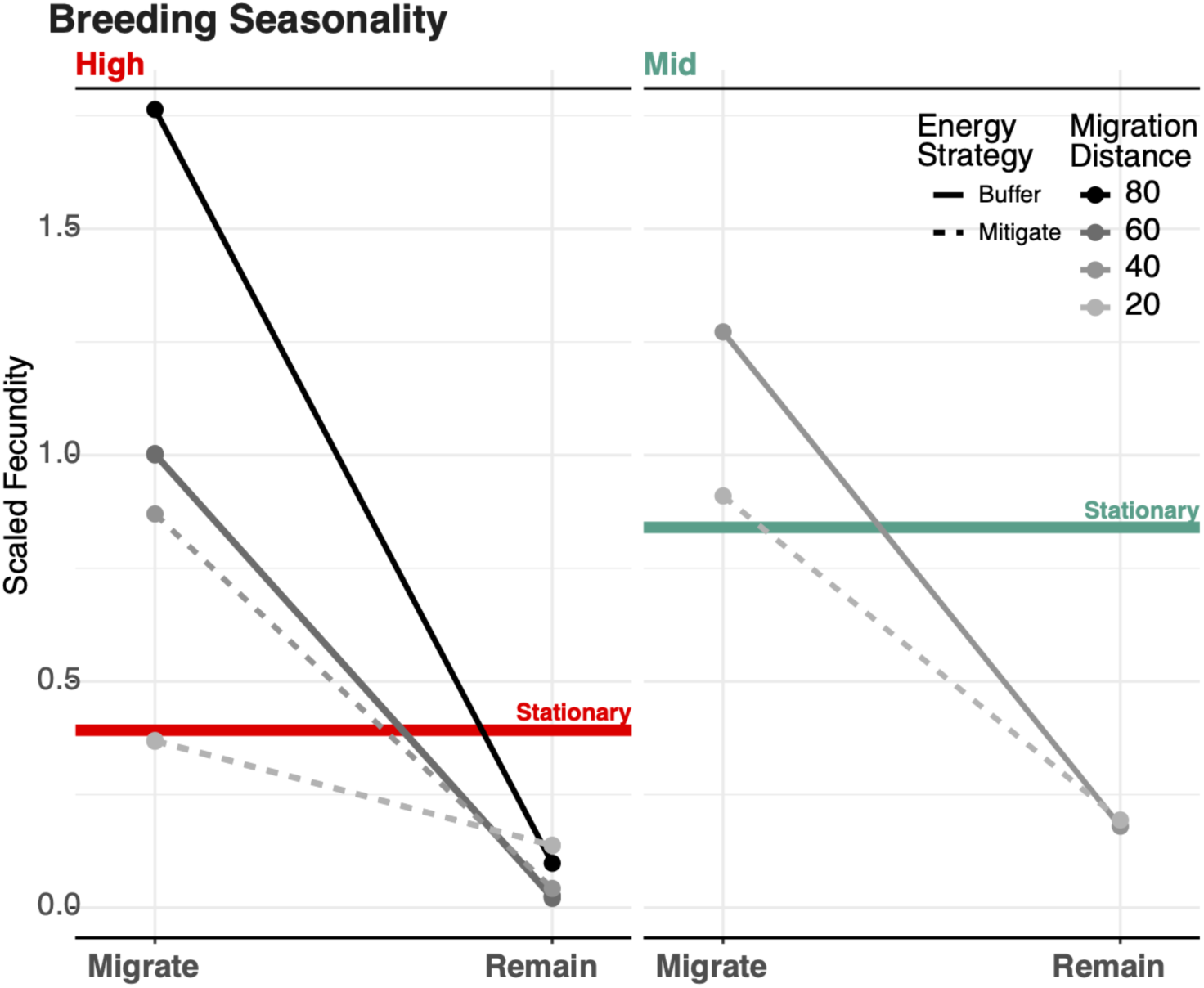
Fecundity of optimized parameterizations of migratory strategies in migratory scenarios compared to simulations in which they do not migrate and remain in the breeding area. Strategies breeding in high seasonality locales in the left panel and mid-seasonal locales on the right with line darkness increasing with migration distance. Solid greyscale lines indicate buffering energy management strategies and mitigation strategies are depicted as dashed lines. Fecundity of stationary strategies depicted as colored lines in each panel for comparison. Values for fecundity are scaled.

### Empirical Analysis

Our macroecological analysis broadly agreed with the general patterns proposed in our theoretical model - life history covaries with biogeography, behavior, and environmental dynamics (Fig. 5). Both migration distance and environmental niche dissimilarity exhibited positive effects on POL (migration distance: avg. marginal effect = 0.002, PD = 99.35%; niche dissimilarity: avg. marginal effect = 0.037, pd = 84.42%) whereas breeding latitude had no apparent influence on POL (avg. marginal effect < 0.001, PD = 58.83%). Variance decomposition showed a strong influence of phylogeny on POL (variance ratio = 0.82).

**Figure 5.**
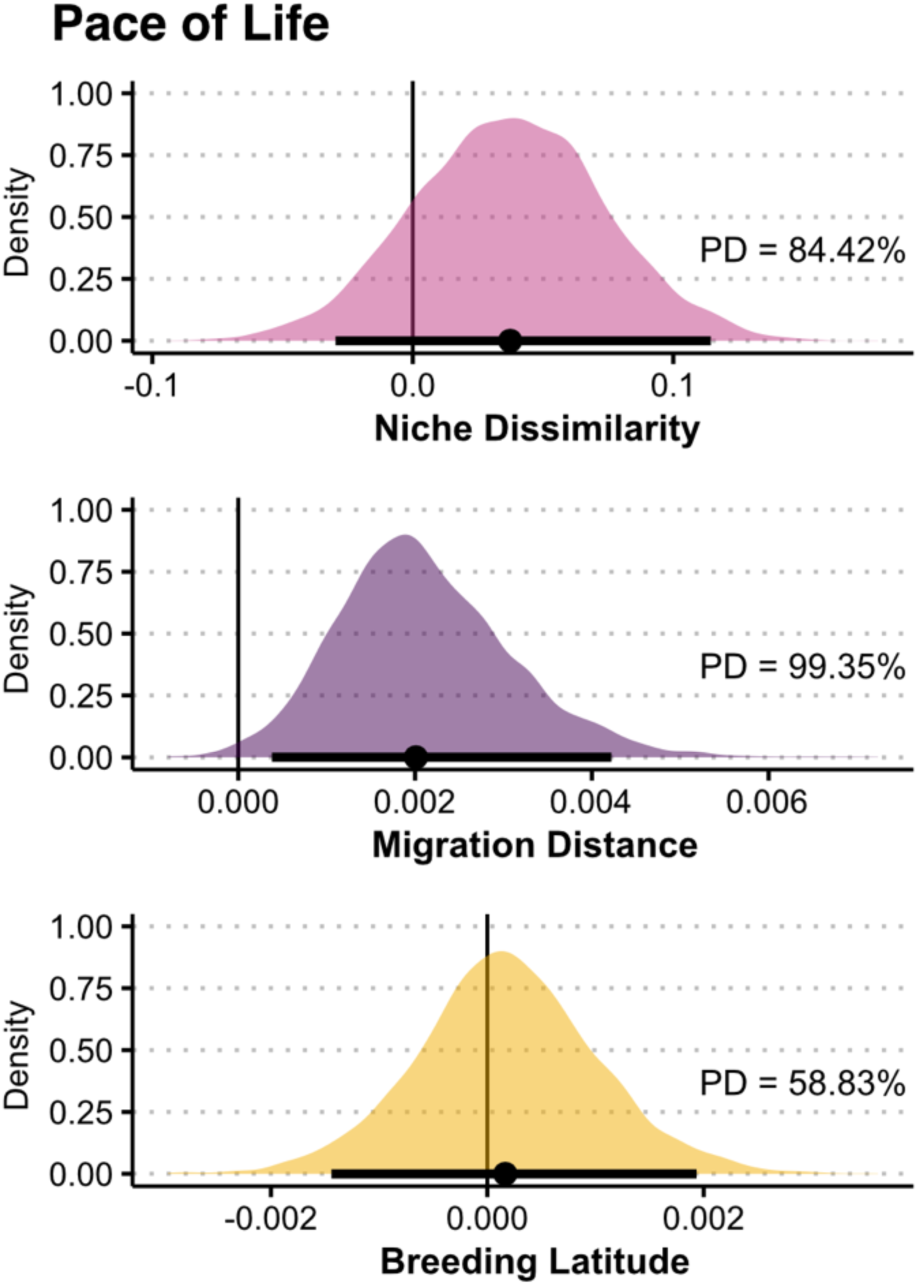
Average posterior marginal effects of niche dissimilarity (top row), migration distance (middle row), and breeding latitude (bottom row; all scaled and centered) on POL based on Bayesian phylogenetic mixed models. POL is measured as a size-normalized estimate of relative maximum somatic mobilization; (see methods; small values indicate fast POL and vice versa). The Bayesian probability of direction (PD) for each marginal effect is displayed at the bottom right of each panel.

## Discussion

### Migration integrates biogeography, behavior, and energetics

Our model suggests a complex relationship between behavior, environment, and life history is fundamentally mediated by metabolism. We observed multiple distinct viable life history strategies within any given migratory strategy (Fig 2A). Some of this diversity in strategy clearly arises from the interaction of behavioral modes and biogeographic factors which are also relevant to life history. For example, the three stationary strategies we considered showed markedly different life histories despite shared behavior (Fig 3). Consistent with previous reports, high energy availability and steep seasonal dynamics at high latitudes push organisms towards fast POL strategies (Mueller & Diamond 2001; Wiersma et al. 2007b; Halali et al. 2020; Yanco et al. 2022). Our results extend these findings to show that behavioral manipulation of the realized seasonal dynamics (via movement) can further modify energetic strategies by interacting with developmental constraints and affecting net energy availability.

Migratory phenology emerged as a fundamental axis of life history variation, primarily trading off the timing and speed of migration with energy allocation strategies. Previous work in boreal breeding birds found that migration distance (a proxy for breeding season duration) negatively covaried with clutch size and annual fecundity such that species with the longest breeding seasons were able to increase the number of young produced (Winger & Pegan 2021). Similarly, we found greater investment in self in strategies that maximized time on migration and non-breeding grounds. This result could relate to previous reports showing, counterintuitively, longer migrations were associated with better body condition (Ortega 2023) and higher annual survival (Zuniga et al 2017) in partially migratory systems. Finally, the apparent importance of phenology suggests that incorporating variable migration speeds could be especially fruitful extensions of the model.

We found that energy management strategy was related to but distinct from migration distance alone. Together, these factors explained substantial variation in optimized life history parameters (Fig 3A) and acted primarily via differential modulation of migration and developmental timing, and energy allocation. Longer migrations tended to result in more buffered energy strategies (by design) but did not perfectly covary, highlighting the utility of using energetics as the common currency to parse behavior-life history relationships.

Our theoretical results suggest that migration mediates metabolic and temporal trade-offs by altering net energy availability (Fig 2.) but differentially based on energy management strategy and migration distance (Fig. 3). Energy-buffering strategies exhibited slower POL than mitigating and coping strategies (Fig. 3A); Short-distance strategies exhibited faster POL than other migratory strategies (Fig. 3B). Similarly, relative investment in self, at the expense of reproduction, was driven by a combination of niche dissimilarity (roughly analogous to energy management strategy) and migration distance in 265 species of migrant birds. Slower POL was associated with greater migration distance but with greater rather than less niche dissimilarity.

However, the metric only measures differences in niche between endpoints and not across the annual cycle, which might explain the converse result. Additionally, some of the environmental variables used in the dissimilarity metric may be unrelated to energy availability. Despite this, empirical results were congruent with the general theoretical prediction that variation in POL is driven by a combination of migratory distance and differences in endpoint seasonality.

### Implications for the evolution of avian migration

Our finding that multiple combinations of life history parameters are theoretically conducive to a migratory lifestyle sheds new light on the apparent facility with which birds can switch between migratory and sedentary lifestyles. Among birds, migration can emerge and disappear over very short timescales (Berthold *et al*. 1992; Garcia-Perez *et al*. 2013). Indeed, phylogenetic analyses suggest that such transitions between migratory states have occurred so frequently over the evolutionary history of class Aves that the potential to migrate is widely viewed as basal to the clade, requiring only minor phenotypic changes to switch states (Zink 2011; Zink & Gardner 2017). In addition to modifying behavioral, morphological, and physiological traits related to migration, any population in the process of switching between migratory states must also move toward a metabolic strategy that maximizes evolutionary fitness. The expected distance in metabolic trait space between a resident life history strategy and a migratory strategy would be lower if, as our results suggest, there are multiple potential metabolic arrangements amenable to a migratory lifestyle. In other words, transitioning from migrant to non-migrant (or vice versa) may not require a radical metabolic reorganization. Thus, with respect to life history, it is perhaps unsurprising that migration has emerged multiple times because there are so many putatively viable migratory life histories which are only partially separable from entirely non-migratory life histories.

Relatedly, our *in silico* experiment showed the steepest declines in reproductive potential when restricting high latitude migrants to a resident strategy. The net energy benefit of migration increased with seasonality of the breeding location and migratory distance, implying that the emergence of a long-distance migration from high latitude residents is associated with a large increase in reproductive potential that far exceeds the best-performing stationary solution. Recent work has suggested a high-latitude origin for most avian migrations as opposed to earlier theories which posit a “southern home” (Bruderer & Salewski 2008; Winger *et al*. 2011, 2019; Yanco et al. 2021). Our results provide theoretical justification for these observations in showing large realized fitness benefits of migration as compared to otherwise optimized non-migratory strategies.

### Model assumptions and extensions

While our model accounts for several elements of dynamic environments and labile behaviors, it is not a comprehensive account of all factors potentially influencing life history optimization. The current instantiation of our model focuses on intrinsically controlled vital rates in concert with environmental dynamism and behavioral plasticity. Future work could extend our model to account for any number of additional factors, including especially other potential extrinsic drivers or phenomena that transpire over shorter timescales. For example, in our model individuals acquire energy at the same rate during migratory movements as while stationary.

Forcing dynamic activity budgeting at a sub-daily resolution (rather than assuming constant foraging rates and efficiencies regardless of migration activity) could affect optimized values of *v̇*. We hypothesize that reduced foraging efficiency during migration might result in solutions showing slower mobilization and/or larger reserve capacity for migratory strategies rather than the faster mobilizations that we observed. Additionally, our simple movement model assumes constant migration speed and symmetry in duration of migrations to and from breeding locations. A more complex movement model could investigate effects of stepwise migration and finer scale speed regulation on trade-off dynamics.

We also did not incorporate extrinsic sources of mortality which have the potential to strongly influence fitness of otherwise viable and fecund life history strategies. Extrinsic mortality during migration, while traditionally difficult to estimate (Rushing 2019), is potentially substantial (Rushing *et al*. 2016; Ward *et al*. 2018; Buechley *et al*. 2021). Similarly, juvenile survival and recruitment is a notoriously important phenomenon for understanding population vital rates (Charnov & Schaffer 1973) for which our model does not account. Finally, differential predation between winter and breeding areas is thought to be an important factor influencing the fitness benefit of migration (Martin 2015). Model extensions that stochastically integrate extrinsic mortality, especially environmentally mediated mortality, would shed light on how animals may compensate for extrinsic mortality via energetic buffering (i.e., aiming for “excess” intrinsic survival rates to counter unpredictable extrinsic mortality).

In addition to energetics in the form of food, ambient temperatures are thought to play an important role in shaping migrations (Corkeron & Connor 1999; Wikelski *et al*. 2003b; Wiersma *et al*. 2007a; Clerc & McGuire 2021). Our model assumes that seasonal temperature variation is reflected in seasonal availability of energy and acquisition rates which scale with organismal surface area. More complex thermoregulatory dynamics would allow exploration of how energy acquisition from the environment relates to the costs of maintaining homeostasis in that environment. For example, accounting for trade-offs between metabolic costs associated with molt/maintenance of body insulation and movement performance could shed light on interactions between molting phenology, migratory strategy, and life history.

Furthermore, even though we sought to maintain model generality, our IBM simulation assumed oviparity, first year reproduction, and initiation of migration at maturation; all of which are generally most consistent with avian life histories. Alternate IBM specifications of reproductive timing, simulation length, movement constraints, or metabolism would allow for more nuanced investigation into developmental trade-off dynamics in taxa that deviate from these constraints. For example, the standard DEB model can be modified to account for fetal development to examine gestational trade-off dynamics in mammals or to account for multiple life stages to examine life-cycle dynamics in invertebrates.

## Conclusion

Our work shows that rather than a singular response to environment, seasonal migrations are actually a diverse set of adaptive strategies that integrate reciprocal relationships between dynamic and spatially-varying environments, plastic behavioral adaptations, and apparently labile metabolic organizations, all of which jointly produce fitness (Hutchinson 1948; Piersma & van Gils 2019). Viewed as such, our mechanistic model provides novel theoretical insights into the emergence of migration across taxa. Moreover, our results clearly demonstrate that behaviors - which both cost energy and modify environmental context - are integral components of life history strategy, providing mathematical support to the Extended POL Theory (Dammhahn *et al*. 2018). Finally, our work shows the promise of approaches that simultaneously consider these reciprocally related aspects of organismal ecology (behavior, environment, and life history) and the unifying power of individual energetics as common currency to relate the three.

## Acknowledgements

We thank Bas Kooijman whose correspondence about our previous work helped to inform aspects of this study. AKP was supported by the Fritz L. Knopf Fellowship in Avian Conservation. SWY was partially supported by the Max Planck – Yale Center for Biodiversity Movement and Global Change.

## Literature Cited

Abrahms, B., Aikens, E.O., Armstrong, J.B., Deacy, W.W., Kauffman, M.J. & Merkle, J.A. (2021). Emerging Perspectives on Resource Tracking and Animal Movement Ecology. Trends Ecol. Evol., 36, 308–320.

Alerstam, T., Bäckman, J., Grönroos, J., Olofsson, P. & Strandberg, R. (2019). Hypotheses and tracking results about the longest migration: The case of the arctic tern. Ecol. Evol., 9, 9511–9531.

Alves, J.A., Gunnarsson, T.G., Hayhow, D.B., Appleton, G.F., Potts, P.M., Sutherland, W.J., et al. (2013). Costs, benefits, and fitness consequences of different migratory strategies. Ecology, 94, 11–17.

Augustine, S., Lika, K., & Kooijman, S. A. (2019). Altricial-precocial spectra in animal kingdom. Journal of Sea Research, 143, 27–34.

Berthold, P., Helbig, A.J., Mohr, G. & Querner, U. (1992). Rapid microevolution of migratory behaviour in a wild bird species. Nature, 360, 360668a0.

BirdLife International and Handbook of the Birds of the World. (2016). Bird species distribution maps of the world.

Brown, J.M., Bouten, W., Camphuysen, K.C.J., Nolet, B.A. & Shamoun-Baranes, J. (2023). Energetic and behavioral consequences of migration: an empirical evaluation in the context of the full annual cycle. Sci. Rep., 13, 1210.

Bruderer, B. & Salewski, V. (2008). Evolution of bird migration in a biogeographical context. J. Biogeogr., 35, 1951–1959.

Buechley, E.R., Oppel, S., Efrat, R., Phipps, W.L., Carbonell Alanís, I., Álvarez, E., et al. (2021). Differential survival throughout the full annual cycle of a migratory bird presents a life-history trade-off. J. Anim. Ecol.

Bürkner, P.-C. (2017). brms: An R Package for Bayesian Multilevel Models Using Stan. Journal of Statistical Software.

Campos-Candela, A., Palmer, M., Balle, S., Álvarez, A. & Alós, J. (2019). A mechanistic theory of personality-dependent movement behaviour based on dynamic energy budgets. Ecol. Lett., 22, 213–232.

Charnov, E.L. & Schaffer, W.M. (1973). Life-History Consequences of Natural Selection: Cole’s Result Revisited. Am. Nat., 107, 791–793.

Clerc, J. & McGuire, L.P. (2021). Considerations of varied thermoregulatory expressions in migration theory. Oikos, 130, 1739–1749.

Cody, M.L. (1966). A GENERAL THEORY OF CLUTCH SIZE. Evolution, 20, 174–184.

Cohen, J. & Jetz, W. (2023). Diverse strategies for tracking seasonal environmental niches at hemispheric scale. Global Ecology and Biogeography 32.9 (2023): 1549–1560.

Corkeron, P.J. & Connor, R.C. (1999). Why do baleen whales migrate?1. Mar. Mamm. Sci., 15, 1228–1245.

Dammhahn, M., Dingemanse, N.J., Niemelä, P.T. & Réale, D. (2018). Pace-of-life syndromes: a framework for the adaptive integration of behaviour, physiology and life history. Behav. Ecol. Sociobiol., 72, 62.

Dubuc-Messier, G., Réale, D., Perret, P. & Charmantier, A. (2017). Environmental heterogeneity and population differences in blue tits personality traits. Behav. Ecol., 28, 448–459.

Garamszegi, L.Z. (Ed.). (2014). Modern Phylogenetic Comparative Methods and Their Application in Evolutionary Biology: Concepts and Practice. Springer, Berlin, Heidelberg.

Garcia-Perez, B., Hobson, K.A., Powell, R.L., Still, C.J. & Huber, G.H. (2013). Switching hemispheres: a new migration strategy for the disjunct Argentinean breeding population of Barn Swallow (Hirundo rustica). PLoS One, 8, e55654.

Gebhardt-Henrich, S. & Richner, H. (1998). Causes of growth variation and its consequences for fitness. Oxford Ornithology Series, 8, 324–339.

Ghalambor, C.K. & Martin, T.E. (2001). Fecundity-survival trade-offs and parental risk-taking in birds. Science, 292, 494–497.

Gómez, C., Tenorio, E., Montoya, P. & Cadena, C. (2016). Niche-tracking migrants and niche-switching residents: evolution of climatic niches in New World warblers (Parulidae). Proc R Soc B, 283, 20152458.

Halali, S., van Bergen, E., Breuker, C.J., Brakefield, P.M. & Brattström, O. (2020). Seasonal environments drive convergent evolution of a faster pace-of-life in tropical butterflies. Ecol. Lett.

Hamblin, S. (2013). On the practical usage of genetic algorithms in ecology and evolution. Methods in Ecology and Evolution, 4(2), 184–194.

Harvey, P.H. & Zammuto, R.M. (1985). Patterns of mortality and age at first reproduction in natural populations of mammals. Nature, 315, 319–320.

Hutchinson, G.E. (1948). Circular causal systems in ecology. Ann. N. Y. Acad. Sci., 50, 221–246.

Jetz, W., Sekercioglu, C.H. & Böhning-Gaese, K. (2008). The worldwide variation in avian clutch size across species and space. PLoS Biol., 6, 2650–2657.

Jetz, W., Thomas, G.H., Joy, J.B., Hartmann, K. & Mooers, A.O. (2012). The global diversity of birds in space and time. Nature, 491, 444–448.

Kooijman, S.A.L.M. (2010). Dynamic Energy Budget Theory for Metabolic Organisation. Cambridge University Press.

Lack, D. (1947). The significance of clutch-size. Ibis, 89, 302–352.

Laskowski, K.L., Moiron, M. & Niemelä, P.T. (2021). Integrating Behavior in Life-History Theory: Allocation versus Acquisition? Trends Ecol. Evol., 36, 132–138.

Levins, R. (1962). Theory of Fitness in a Heterogeneous Environment. I. The Fitness Set and Adaptive Function. Am. Nat., 96, 361–373.

Marques, G.M., Augustine, S., Lika, K., Pecquerie, L., Domingos, T. & Kooijman, S.A.L.M. (2018). The AmP project: Comparing species on the basis of dynamic energy budget parameters. PLoS Comput. Biol., 14, e1006100.

Martin, T.E. (1996). Life History Evolution in Tropical and South Temperate Birds: What Do We Really Know? J. Avian Biol., 27, 263–272.

Martin, T.E. (2015). Age-related mortality explains life history strategies of tropical and temperate songbirds. Science, 349, 966–970.

Merkle, J.A., Monteith, K.L., Aikens, E.O., Hayes, M.M., Hersey, K.R., Middleton, A.D., et al. (2016). Large herbivores surf waves of green-up during spring. Proc. Biol. Sci., 283, 20160456.

Lynch. M. (1991). Methods for the Analysis of Comparative Data in Evolutionary Biology. Evolution, 45, 1065–1080.

Moreau, R.E. (1944). Clutch-size: a comparative study, with special reference to African birds. Ibis, 86, 286–347.

Mueller, P. & Diamond, J. (2001). Metabolic rate and environmental productivity: well-provisioned animals evolved to run and idle fast. Proc. Natl. Acad. Sci. U. S. A., 98, 12550–12554.

Muller, E.B. & Nisbet, R.M. (2000). Survival and production in variable resource environments. Bull. Math. Biol., 62, 1163–1189.

Nakazawa, Y., Peterson, A.T., Martínez-Meyer, E. & Navarro-Sigüenza, A.G. (2004). Seasonal Niches of Nearctic-Neotropical Migratory Birds: Implications for the Evolution of Migration. Auk, 121, 610–618.

van Noordwijk, A.J. & de Jong, G. (1986). Acquisition and Allocation of Resources: Their Influence on Variation in Life History Tactics. Am. Nat., 128, 137–142.

Ortega, A. C. (2023). Evaluating Resource Tracking and the Benefit of Migration for a Temperate Ungulate (Doctoral dissertation, University of Wyoming).

Piersma, T. & van Gils, J.A. (2019). The Flexible Phenotype: A Body-Centred Integration of Ecology, Physiology, and Behaviour. Oxford University Press, New York, NY, USA.

R Core Team (2022). R: A language and environment for statistical computing. R Foundation for Statistical Computing, Vienna, Austria.

Réale, D., Dingemanse, N.J., Kazem, A.J.N. & Wright, J. (2010). Evolutionary and ecological approaches to the study of personality. Philos. Trans. R. Soc. Lond. B Biol. Sci., 365, 3937–3946.

Ricklefs, R.E. (2000). Density Dependence, Evolutionary Optimization, and the Diversification of Avian Life Histories. Condor, 102, 9–22.

Ricklefs, R.E. & Wikelski, M. (2002). The physiology/life-history nexus. Trends Ecol. Evol., 17, 462–468.

Rushing, C.S. (2019). Estimability of migration survival rates from integrated breeding and winter capture–recapture data. Ecol. Evol.

Rushing, C.S., Hostetler, J.A., Sillett, T.S., Marra, P.P., Rotenberg, J.A. & Ryder, T.B. (2017). Spatial and temporal drivers of avian population dynamics across the annual cycle. Ecology, 98, 2837–2850.

Rushing, C.S., Ryder, T.B. & Marra, P.P. (2016). Quantifying drivers of population dynamics for a migratory bird throughout the annual cycle. Proc. Biol. Sci., 283.

Sæther, B.-E. (1988). Pattern of covariation between life-history traits of European birds. Nature.

Sillett, S.T. & Holmes, R.T. (2002). Variation in survivorship of a migratory songbird throughout its annual cycle. J. Anim. Ecol., 71, 296–308.

Soriano-Redondo, A., Gutiérrez, J.S., Hodgson, D. & Bearhop, S. (2020). Migrant birds and mammals live faster than residents. Nat. Commun., 11, 5719.

Spiegel, O., Leu, S.T., Sih, A., Godfrey, S.S. & Bull, C.M. (2015). When the going gets tough: behavioural type-dependent space use in the sleepy lizard changes as the season dries. Proc. Biol. Sci., 282.

Stearns, S.C. (1989). Trade-Offs in Life-History Evolution. Funct. Ecol., 3, 259–268.

Stearns, S.C. (1992). The Evolution of Life Histories. OUP Oxford.

Thorup, K., Tøttrup, A.P., Willemoes, M., Klaassen, R.H.G., Strandberg, R., Vega, M., et al. (2017). Resource tracking within and across continents in long-distance bird migrants. Sci Adv, 3, e1601360.

Ward, M.P., Benson, T.J., Deppe, J., Zenzal, T.J., Diehl, R.H., Celis-Murillo, A., et al. (2018). Estimating apparent survival of songbirds crossing the Gulf of Mexico during autumn migration. Proceedings of the Royal Society B: Biological Sciences, 285, 20181747.

Wiersma, P., Chappell, M.A. & Williams, J.B. (2007a). Cold- and exercise-induced peak metabolic rates in tropical birds. Proc. Natl. Acad. Sci. U. S. A., 104, 20866–20871.

Wiersma, P., Muñoz-Garcia, A., Walker, A. & Williams, J.B. (2007b). Tropical birds have a slow pace of life. Proc. Natl. Acad. Sci. U. S. A., 104, 9340–9345.

Wikelski, M. & Ricklefs, R.E. (2001). The physiology of life histories. Trends Ecol. Evol., 16, 479–481.

Wikelski, M., Spinney, L., Schelsky, W., Scheuerlein, A. & Gwinner, E. (2003a). Slow pace of life in tropical sedentary birds: a common-garden experiment on four stonechat populations from different latitudes. Proceedings of the Royal Society of London. Series B: Biological Sciences, 270, 2383–2388.

Wikelski, M., Tarlow, E.M., Raim, A., Diehl, R.H., Larkin, R.P. & Henk Visser, G. (2003b). Costs of migration in free-flying songbirds. Nature, 423, 704–704.

Williams, G.C. (1966). Natural Selection, the Costs of Reproduction, and a Refinement of Lack’s Principle. Am. Nat., 100, 687–690.

Winger, B.M., Auteri, G.G., Pegan, T.M. & Weeks, B.C. (2019). A long winter for the Red Queen: rethinking the evolution of seasonal migration. Biol. Rev. Camb. Philos. Soc., 94, 737–752.

Winger, B.M., Lovette, I.J. & Winkler, D.W. (2011). Ancestry and evolution of seasonal migration in the Parulidae. Proc Royal Soc B Biological Sci, 279, 610–618.

Winger, B.M. & Pegan, T.M. (2021). Migration distance is a fundamental axis of the slow-fast continuum of life history in boreal birds. Ornithology.

Yanco, S.W., Linkhart, B.D., Marra, P.P., Mika, M., Ciaglo, M., Carver, A., et al. (2021). Niche dynamics suggest ecological factors influencing migration in an insectivorous owl. Ecology.

Yanco, S.W., Pierce, A.K. & Wunder, M.B. (2022). Life history diversity in terrestrial animals is associated with metabolic response to seasonally fluctuating resources. Ecography, 2022.

Zink, R.M. (2011). The evolution of avian migration. Biol. J. Linn. Soc. Lond., 104, 237–250.

Zink, R.M. & Gardner, A.S. (2017). Glaciation as a migratory switch. Sci Adv, 3, e1603133.

Zúñiga, D., Gager, Y., Kokko, H., Fudickar, A.M., Schmidt, A., Naef-Daenzer, B., et al. (2017). Migration confers winter survival benefits in a partially migratory songbird. Elife, 6.

Zurell, D., Gallien, L., Graham, C.H. & Zimmermann, N.E. (2018). Do long-distance migratory birds track their niche through seasons? J. Biogeogr., 45, 1459–1468.

