## Appendix S1 (Model description) for "Seasonal migration alters energetic trade-off optimization and shapes life history" (from another version of this preprint: version 1, 2023-09-15)

### Appendix S1: Supplemental Model Description

Allison K. Pierce Scott W. Yanco Michael B. Wunder

Sep 12, 2023

### 1 IBM Model Overview, Design, and Detail (ODD)

The structure of this model documentation generally follows ODD protocol [1] to describe the individual-based model description and optimization routine from Pierce et al. "Seasonal migration alters energetic trade-off optimization and shapes life history"

#### 1.1 Purpose

The purpose of this individual-based model (IBM) is to simulate individual migratory movement and metabolism in an environment with seasonally fluctuating energetic resources.

The IBM is comprised of two sub-models:

- A movement model to simulate migration across a landscape of dynamic energy availability.
- An individual model of daily metabolism based on Dynamic Energy Budget (DEB) theory [3] to simulate individual acquisition and allocation of energy.

This model was developed to examine how migratory behavior affects metabolic and temporal trade-off dynamics which shape life-history and pace of life by using a genetic algorithm to identify fitness optimized parameterizations for migratory and stationary movement strategies.

Ultimately, the goal of the model is to provide a theory based hypothesis generation frame-work for increased understanding of how life-history strategy is affected by behaviors which alter metabolic and temporal trade-off dynamics, like migration. IBM simulation and optimization routines are implemented in the R language.

#### 1.2 Entities, state variables, and scales

The IBM consists of two primary entities, a set of parameters representing an individual organism and the environment, an energy landscape consisting of a time dimension and a single spatial dimension. Individual organisms are simulated in isolation (interact only with the environment not with each other).

Individuals are characterized by 4 primary DEB state variables:

1. Structure ($l$), scaled length of an individual (size relative to maximum).
2. Energy reserve density ($e$), relative capacity of intermediate storage of energy between assimilation and mobilization processes (proportion of maximum).
3. Scaled maturation energy ($u_H$), cumulative energetic investment towards maturation (unitless).
4. Scaled reproductive energy buffer ($u_R$), cumulative energetic investment towards reproduction (unitless).

The environment is a simplified virtual landscape of seasonally available energy on earth ($f$) across latitude from pole to pole. At each latitude location, available energy ($f$) varies across 365 discrete time steps (1 year) as a single oscillation of a sine function whose amplitude and phase varies spatially across 180 continuous latitude units such that amplitude is at maximum at the lowest and highest latitude, decreasing towards the mid-point, to reflect decreasing severity of seasonality towards the equator. For locations below the mid-point phase is shifted by half a wavelength relative to those above the mid-point to reflect opposing seasonality between hemispheres.

Table 1.1: State variables

| Symbol | Units | Description and value range |
|---|---|---|
| **Environment** | | |
| step | integer; day | Current time step; [1..365] |
| location | real number, degrees latitude | Current location; $[-90, 90]$ |
| $f$ | real number; - | Proportion of total available energy for current location and time step; $(0, 1]$ |
| **Organism** | | |
| $l$ | real number, - | Structural length scaled relative to maximum length; $(0, 1]$ |
| $e$ | real number, - | Energy reserve density. Reserve energy per unit of structural volume ($l^3$) relative to maximum; $(0, 1]$ |
| $u_H$ | real number, - | Cumulative energy invested into maturity scaled unitless; $(0, +\infty)$ |
| $u_R$ | real number, - | Cumulative energy invested into reproduction scaled unitless; $(0, +\infty)$ |

Table 1.2: Organism/Individual parameters

| Symbol | Units | Description and value range |
|---|---|---|
| $\kappa$ | real number; - | Proportion of mobilized energy invested into somatic processes; $(0, 1)$ |
| $\dot{v}$ | real number; Length/day | energy conductance rate; $(0, 1]$ |
| $m$ | real number; - | scaled locomotion maintenance rate; $[0, 1]$ |
| $l$ | real number, - | Structural length scaled relative to maximum length; $(0, 1]$ |
| $g$ | real number, - | Energy investment ratio; $(0, +\infty)$ |
| $u_H^b$ | real number, - | Scaled maturity at birth; $(0, +\infty)$ |
| $u_H^p$ | real number, - | Scaled maturity at sexual maturity (puberty) ; $(0, +\infty)$ |
| $d_M$ | integer; day | Migration duration; (1..365) |
| $d_{NB}$ | integer; day | Stationary duration at non-breeding location; (1..365) |
| $d_B$ | integer; day | Stationary duration at breeding location; (1..365) |
| $s_H^{bp}$ | real number, - | Maturity ratio; $(0, 1]$ |
| $\dot{k}_M$ | real number, - | Somatic maintenance rate coefficient; $(0, +\infty)$ |
| $a_b$ | integer; day | Age at birth/incubation duration; (1..365) |
| $a_p$ | integer; day | Age at sexual maturity; (1..365) |
| $u_E^0$ | real number, - | Scaled cost of one egg; $(0, +\infty)$ |

#### 1.3 Process overview and scheduling

Process: The IBM simulation evaluates a daily energy acquisition and allocation for an individual over the course of one year (365 days) from birth on day 1, through sexual maturity, until a clutch of eggs is produced and hatch. All individuals are born in an environment at the peak of seasonal availability in energy for their starting location and produce a clutch of eggs at a time such that they develop and are born/hatch when availability peaks again on day 365. The number of offspring produced is based on the amount of cumulative energy invested into reproduction divided by the cost of a egg. Egg cost is based on the individuals condition at production (maternal effect) and the assumption that offspring size and age at birth is the same as the parent (inherited traits). Migratory individuals begin migrating to their non-breeding location when they reach adulthood (sexual maturity) and must also return to their initial breeding location before producing a clutch of eggs.

Schedule: An individuals is born on the first time step in a pre-specified location. After each time step DEB state variables are updated based on numerically solving a set of DEB differential equations (see Section 1.7.1 for detail). If individuals are migratory, at the end of each time step the movement sub-model is invoked to evaluate if individuals move and to what location. At the last time step, solutions are checked for viability and fecundity is calculated for viable solutions. Viable solutions and their corresponding parameters are then processed by the optimization routine.

The following pseudo-code generally describes the scheduling of events in the IBM:

Calculate change in $e$

If not mature

- Calculate change in $u_H$

If mature

- Calculate change in $u_R$

Calculate change in $l$

- If change in $l < 0$
  - Re-calculate change in maturity/reproductive buffer based on starvation rules

Update state variables

- If $e < 0$
  - Die

If migratory, mature, and current time step is not during the non-breeding period

- Move to a new location If current time step is before the non-breeding period
  - Move towards non-breeding location If current time step is after the non-breeding period and current location
  - Move towards breeding location If time step = 365

Calculate fecundity

- If fecundity < 1
  - Die

If migration incomplete

- Die

#### 1.4 Design concepts

##### 1.4.1 Basic principles

This model uses individual level simulations to mechanistically model interactions between seasonality, movement behavior, and metabolism to inform construction of hypotheses about how trade-off dynamics shape life history and pace of life of migratory animals. To evaluate movement and metabolism in a dynamic environment we used an IBM implementation of a dynamic energy budget (DEB) model to numerically solve daily metabolism over the course of 1 year (365 time steps). The DEB framework [3] is particularly suited for this as it models energy acquisition and allocation as a series of metabolic trade-offs downstream of the fundamental life history trade-off between investment into self versus reproduction. We incorporate modeling of additional metabolic and temporal trade-offs associated with migratory behavior by integrating a simple migratory movement model with the DEB and adding a new parameter for migration locomotion cost. The model is formulated such that quantities of energy and size are relative, as we sought to maintain generality. However, as currently implemented, our IBM simulation assumes oviparity, first year reproduction, and maturation prior to migration; all of which are generally most consistent with avian life histories.

##### 1.4.2 Emergence

Combinations of individual traits and migratory behaviors that are viable and allow for reproduction emerge from the properties of metabolic organization in a dynamic environment.

##### 1.4.3 Adaptation

Movement is the only adaptive behavior in the IBM. Individuals move based on their level of maturity and their migration strategy. Migration strategies are defined by parameters for breeding and non-breeding locations, migration speed, and duration of the non-breeding period.

##### 1.4.4 Objectives

The objective measures individuals use to decide to move is their maturity level, the current time step, and their current location. Individuals only move if they are mature, the non-breeding period has not begun, or the non-breeding period is complete and they have not arrived at the breeding location. If the non-breeding period has not begun then individuals move towards non-breeding locations. If the current time is after the non-breeding period individuals move towards breeding areas. Individuals move towards their target location until their location is less than one movement away based on their pre-determine migration speed.

##### 1.4.5 Learning

Individual learning is not modeled.

##### 1.4.6 Prediction

Predictive behaviors of individuals are not modeled.

##### 1.4.7 Sensing

Individuals can sense their current state, location, and time.

##### 1.4.8 Interaction

Individuals do not interact.

##### 1.4.9 Stochasticity

No stochastic processes are modeled in the IBM.

##### 1.4.10 Collectives

None; individuals are simulated in isolation.

##### 1.4.11 Observation

The IBM outputs the living status, final states, parameter values and location of the individual at each time step which can be be further analyzed in R to evaluate model performance. Movement paths can be visualized from the locations. The energy availability over the path can also be obtained using the locations with the lookupf_bylat() function. Metabolic and temporal trade-offs can be observed with a principle component analysis of scaled and centered individual parameter values and visualized with a bi-plot.

#### 1.5 Initialization

Table 1.3: State variables

| Variable | Initial value |
|---|---|
| step | 1 |
| location | Starting location specified in model parameters |
| $f$ | Determined by starting location and environment function. |
| $l$ | $l_b$ |
| $e$ | energy available in the starting location $f$ but later adjusted. (see Section 2) |
| $u_H$ | $u_H^b$ |
| $u_R$ | 0 |

Table 1.4: Global model parameters

| Parameter | Initial value | Description |
|---|---|---|
| worldsize | 180 | Total distance of simulated environment |
| steps | 365 | Simulation length in days |
| meanrange | 0.6,0.5 | Range of mean $f$ across seasonal environment. Intialized with mean $f$ of at mid-point location (0) at 0.6 and linearly decreasing to 0.5 at location 90 and -90. |
| movement | varies - see below | Boolean to enable migratory movement sub-model |
| startloc | varies - see below | Breeding location |
| endloc | varies - see below | Non-breeding location |

Table 1.5: Movement strategy specific initializations

| startloc | endloc | movement |
|---|---|---|
| 80 | -80 | T |
| 80 | -40 | T |
| 80 | 0 | T |
| 80 | 40 | T |
| 40 | -40 | T |
| 40 | 0 | T |
| 80 | 80 | F |
| 40 | 40 | F |
| 0 | 0 | F |

See Section 2.1 for initialization of individual-specific parameters for fitness optimization.

#### 1.6 Input data

No external data was used to drive model processes.

#### 1.7 Submodels

##### 1.7.1 DEB sub-model

The metabolic sub-model is a dynamic mechanistic model based on Dynamic Energy Budget (DEB) theory (Koojiman 2010) that models how individual organisms uptake, use, and dissipate energy over their lifespans and across different life stages based on the principles of conservation of energy and mass, thermodynamics, and stoichiometric constraints.DEB theory posits a unified metabolic framework generalizable across taxa based on first principles as compared to species specific bioenergetic models derived from empirical observation. At the basis of this framework is the standard DEB model which is a simplified model that describes the energy fluxes of an organism that does not change shape, feeds on a single food source, and is comprised of two main energetic components – reserves and bodily structure which do not change in chemical composition. Reserves represent energy that has been assimilated by the organism that is available to be used for metabolic processes including growth, maintenance and reproduction. Bodily structure requires energetic maintenance proportional to its volume (and surface area in endotherms) and dictates assimilation of energy proportional to its surface area. This basic model can be extended to accommodate modeling of taxa specific traits including but not limited to changes in body shape, existence of multiple reserve or structure types with different compositions, reproductive buffer handling rules, or varying number of life stages.

In the standard model, organisms in juvenile and adult stages acquire energy from food (uptake) and only a portion of this will be assimilated into reserves with the remainder being excreted as feces (defection). Embryos do not feed from the environment and rely on energy in their reserves until birth. Energy is then mobilized from reserve and allocated at a fixed proportion ($\kappa$) to cover energetic requirements of growth and maintenance with the remainder diverted to maturation and reproduction ($\kappa$). Energy allocated to the soma used for somatic maintenance is used for metabolic work and dissipated to the environment. The remainder is used for growth and fixed into structure. A portion of energy allocated to reproduction/maturation is used for metabolic work to maintain reproductive condition and bodily defense systems (such as the immune system). If the organism is in the adult stage the remainder is accumulated into a reproductive buffer to be converted into gametes. In embryonic and juvenile stages, it is used for maturation processes not associated with increase in body size (growth) such as development of new organ systems or regulation systems. Life stage transitions occur at individual specific thresholds of cumulative energy invested into maturation.

###### 1.7.1.1 Model modification for migration movement

In the standard DEB model, locomotion costs are not explicitly defined but are assumed to come from the somatic maintenance costs when they are independent from foraging rate. We modified the standard model to isolate migration movement costs from somatic maintenance costs via the compound parameter $m$, which is the ratio between metabolic cost of migration locomotion to travel one body length and non-migration somatic maintenance per unit of structural volume. Somatic maintenance costs reflect not only energy spent on cell turnover but also average daily energy expenditure lost to dissipation and not fixed into structure (growth). Here we modified the standard model to differentiate migration movement costs from somatic maintenance costs with a new compound parameter $m$, a ratio between maintenance rate of migration locomotion only and non-migration somatic maintenance. This cost scales with surface area and introduces a new energy rate parameter specific to migratory locomotion $\dot{p}_L$. We adjusted other rate questions to account for this new rate to maintain model structure.

###### 1.7.1.2 Model generalization

In the standard model, a set of 12 individual specific parameters describe the rates or fluxes of energy flow as well as the capacity of the reserve and structure variables and life stage transitions. To characterize dynamics of metabolic processes generically we implemented unitless version of the model (Koojiman 2010) that describes dynamics independent of specific values of energy (Joules) or structural volume (length/mass). Thus, individual specific parameters are scaled by thier maximums or represented in compound parameters (ratios between individual-specific parameters). The DEB model and its derivation is described in detail in Koojiman (2010) and Sousa et al. (2013).

###### 1.7.1.3 DEB IBM parameters

###### Environmental variables

$f$ Scaled functional response $\frac{X}{(K+X)}$

- A Holling type II functional response of the density of food in the environment ($X$) relative to the food density that results in half the maximum food uptake rate ($K$). In our implementation $K$ is constant which assumes all organisms of the same size ingest and search at the same rate in our simulation thus $f$ is a property of the environment as any changes in $f$ are due to changes in food density ($X$).
- Generated from a wave function that emulates a latitudinal gradient of seasonality. See Equation 1.13

##### Individual specific state variables

$e$ Scaled energy reserve density $\frac{[E]}{[E_M]}$

- Proportion of current energy in reserve per unit structural volume ($[E]$) over maximum energy per structural unit ($[E_M]$).

$l$ Scaled structural length $\frac{[L]}{[L_M]}$

- Proportion of current structural length over maximum length.

$u_R$ Scaled cumulative energy invested into reproduction (Reproductive buffer).

$u_H$ Scaled cumulative energy invested into maturation.

##### Individual specific parameters

$\kappa$ Proportion of mobilized reserve energy density committed to somatic maintenance and growth. Higher values mean more energy is allocated towards somatic maintenance and growth.

$\dot{k}_M$ Somatic maintenance rate coefficient. Ratio between somatic maintenance rate per unit of volume $[\dot{p}_M]$ and volume-specific growth costs of structure $[E_G]$. Higher values imply higher maintenance costs or lower growth costs. $\frac{[\dot{p}_M]}{[E_G]}$

$\dot{k}_J$ Maturity maintenance rate coefficient. Analogous to $\dot{k}_M$ with respect to maturity. Ratio between the cost to maintain maturity relative to the cost to mature. To simplify stage transition dynamics, we assume $\dot{k}_J = \dot{k}_M$.

$\dot{v}$ Energy conductance rate. Ratio between surface area linked maximum assimilation rate $\dot{p}_{Am}$ and maximum reserve density $[E_M]$. Controls mobilization rate; higher values mean faster mobilization of energy from reserves per time step. Lower values imply smaller maximum reserve capacity or higher maximum assimilation rates. $\frac{\{\dot{p}_{Am}\}}{[E_M]}$

$g$ Energy investment ratio. Compound parameter that is a ratio of the cost to grow 1 unit of structural volume $[E_G]$ relative to maximum available reserve energy density for growth and maintenance $\kappa[E_M]$. Higher values imply higher growth costs or larger reserve capacity available for somatic branch. $\frac{[E_G]}{\kappa[E_M]}$ Is also equivalent to $\frac{\dot{v}}{\dot{k}_M[L_M]}$

$l_b$ Scaled structural length at birth $\frac{[L]}{[L_M]}$. Proportion of current structural length over maximum length at birth.

$u_H^b$ Scaled maturity at birth. Cumulative energy investment into maturity at time of birth. Calculated from $l_b$ given $u_H^b = (1 - \kappa) * l_b^3$ which is based on the assumptions $\dot{k}_J = \dot{k}_M$, a thermo-neutral environment, and transition to the juvenile state occurs at a fixed structural volume ($\dot{k}_J = \dot{k}_M$).

$u_H^p$ Scaled maturity at sexual maturity (puberty). Cumulative energy investment into maturity at transition to adulthood.

$m$ Cost to move one volume of structure one unit of body length relative to somatic maintenance rate of one unit of volume. $\frac{\dot{p}_L}{\dot{p}_M}$

##### Energy fluxes

$\dot{p}_A$ Assimilation. Flux of assimilated energy from the environment to the reserve energy density per unit of time. Because all individuals are assumed to have the same ingestion rates (and thus the energy loss to defecation) assimilation rates only vary by scaled surface area.

$$\dot{p}_a = fl^2 \tag{1.1}$$

$\dot{p}_M$ Somatic maintenance rate. Flux of mobilized reserve energy density allocated to soma ($\kappa$) used for maintenance processes not related to long scale migratory movements.* assumes ectothermy or endothermy in a thermoneutral environment

$$\dot{p}_M = \kappa l^3 \tag{1.2}$$

$\dot{p}_L$ Somatic maintenance rate of migratory locomotion. Flux of mobilized reserve energy density allocated to soma ($\kappa$) used for maintenance processes related to a migratory movement of $d$ distance units.

$$\dot{p}_L = \kappa l^2 * m * d \tag{1.3}$$

- Migration locomotory costs are assumed to scale with surface area. For simplicity, we also assume that costs scale linearly with distance.

$\dot{p}_S$ Somatic maintenance. Flux of mobilized reserve energy density allocated to soma ($\kappa$) used for all maintenance processes (energy lost to dissapation).

$$\dot{p}_S = \dot{p}_M + \dot{p}_L \tag{1.4}$$

$\dot{p}_G$ Growth. Flux of mobilized reserve energy density allocated to soma ($\kappa$) fixed into structure ($l$).

$$\dot{p}_G = \kappa l^2 \frac{e - l - (m * d)}{1 + e/g} \tag{1.5}$$

$\dot{p}_R$ Reproduction. Flux of mobilized reserve energy density allocated maturation prior to adulthood and reproduction after adulthood.

$$\dot{p}_R = (1 - \kappa) e l^2 \frac{g + : l + (m * d)}{g + e} - u_H \tag{1.6}$$

$\dot{p}_J$ Maturity maintenence. Flux of mobilized reserve energy density allocated maintenance of maturity. Once adult stage is reached $u_H = u_H^P$.

$$\dot{p}_J = u_H \tag{1.7}$$

$\dot{p}_c$ Mobilization. Total flux of mobilized reserve energy density to somatic and reproductive processes per unit of time. Represents the total of all energy fluxes from the reserve.

$$\dot{p}_c = el^2 \frac{g + l + (m * d)}{g + e} = \dot{p}_S + \dot{p}_G + \dot{p}_J + \dot{p}_R \tag{1.8}$$

All fluxes are scaled by $\dot{p}_{Am}[L_M]^2$ to keep dynamics unit-less in length and energy.

###### 1.7.1.4 Model dynamics

Mobilization flux is the sum of all somatic and reproductive fluxes.

$$\kappa \dot{p}_c = \dot{p}_S + \dot{p}_G$$

$$(1 - \kappa) * \dot{p}_c = \dot{p}_J + \dot{p}_R \tag{1.9}$$

Therefore, change in reserve energy density is the difference between assimilation and mobilization. Where mobilization is the sum of all somatic and reproductive fluxes and the sum of the somatic fluxes is the product of mobilization and $\kappa$.

$$de = \frac{g\dot{k}_M}{l^3}(\dot{p}_a - \dot{p}_c - \frac{e}{g\kappa}\dot{p}_g) \tag{1.10}$$

$$du_R = \begin{cases} \dot{k}_M * (1 - \kappa)el^2 \frac{g+l+(m*d)}{g+e} - u_H^P, & \text{if } u_H = u_H^P \\ 0, & \text{if } u_H < u_H^P \end{cases} \tag{1.11}$$

$$du_H = \begin{cases} \dot{k}_M * (1 - \kappa)el^2 \frac{g+l}{g+e} - u_H, & \text{if } u_H < u_H^P \\ 0, & \text{if } u_H = u_H^P \end{cases} \tag{1.12}$$

Energy flux to scaled reserves are re-scaled by $\frac{g\dot{k}_M}{l^3}$ to scale energy relative to maximum energy density $[E_M]$. The term $\frac{e}{g\kappa}\dot{p}_g$ in the $\delta e$ adjusts energy density for growth related changes in structure. Energy flux to scaled maturity/reproductive buffer is scaled by $\dot{k}_M$ to match scaling of maturation thresholds and egg cost.

Energy available in the environment $f$ is calculated as a function of time step $t$ and location $loc$.

$$f = a_{loc} * cos(2 * pi * t/365) + \mu_{f,loc} \tag{1.13}$$

$$a_{loc} = loc/90 * (1 - \mu_{f,loc}) \tag{1.14}$$

$$\mu_{f,loc} = \mu_{f,0} - ((\mu_{f,0} - \mu_{f,90}) * |loc|/90)) \tag{1.15}$$

$$\mu_{f,0} \geq \mu_{f,90} \text{ and } \mu_{f,90} = \mu_{f,-90} \tag{1.16}$$

Amplitude of the wave function ($a$) is highest at $t = 1$ for latitudes $> 0$ and at $t = 365$ for latitudes $< 0$. If mean $f$ ($\mu_f$) over time varies by latitude then $\mu_f$ at the equator is assumed to linearly decrease towards the poles.

##### Starvation dynamics

In starvation conditions, assimilation is less than mobilization and energy is depleted from the reserves (-$de$). Energy is prioritized in order by somatic maintenance, growth, then maturation/reproduction therefore, no energy is allocated to reproduction when reserve energy density ($e$) equals total somatic costs ($\kappa\dot{p}_c$) and zero growth occurs once reserve energy density ($e$) is equal to scaled length ($l$). Once reserves are no longer sufficient for somatic maintenance death follows. We do not allow length ($l$) or maturation level ($u_H$) to shrink or for energy previously allocated to the reproductive buffer $u_R$ to be used.

#### 1.7.2 Movement sub-model

Migratory movement decisions are based on a simplified model of avian migration that assumes individuals migrate when reaching adulthood to a single non-breeding location where they remain stationary for a period of time before returning to the same location they were born to breed. We assume migrations are non-stop and occur at a constant speed. We also assume that migration speeds to and from breeding locations are equal. Although in reality migrations can be much more complex, this simplified model still allows us to many examine behavior mediated trade offs while maintaining a tractable number of parameters for later optimization which is computationally intensive.

###### 1.7.2.1 Movement model Parameters

$s_M$: Migration speed

$d_{NB}$: Duration of non-breeding period in days

$t_{NB}$: Start of non-breeding period

$d$: Distance traveled

$loc$: Location of individual

###### 1.7.2.2 Movement model dynamics

After each time step individuals move towards non-breeding locations at their pre-determined migration speed once sexually mature or depart to breeding locations if duration at non-breeding location has been reached. After departure, individuals stop moving once their location is less than one movement away from their target breeding location.

$$d = \begin{cases} 0, & \text{if } u_H < u_H^P \\ 0, & \text{if } 0 < t_{NB} \leq t \leq t_{NB} + d_{NB} \\ 0, & \text{if } t > t_{NB} + d_{NB} > d_{NB} \text{ and } |loc - loc_B| < s_M, \\ s_M, & \text{otherwise} \end{cases}$$

> **Note**
>
> Once the individual arrives to a location less than one movement away from the non-breeding location ($|loc - loc_B| < s_M$) the start of the non-breeding period $t_{NB}$ is set to the current time step $t$. Until then $t_{NB} = 0$.

$$dloc = \begin{cases} -d, & \text{if } t < t_{NB}, \\ d & \text{otherwise} \end{cases}$$

> **Note**
>
> As written, this assumes the breeding location is north of the non-breeding location ($loc_B > loc_{NB}$) which is true of all of the movement strategies we modeled, if this was not true the conditionals for $\delta loc$ switch. This is accounted for in the model code.

### 2 IBM Parameter Optimization

Heuristic optimization algorithms are useful for efficiently solving multidimensional optimization problems that are analytically intractable and too computationally intensive to solve numerically by brute force [2]. We developed a genetic algorithm to identify fitness-optimized parameters of the IBM for each movement strategy. A genetic algorithm is a heuristic optimization of an objective function (represented here as an IBM) that uses iterative search techniques inspired by genetic mechanisms of natural selection such as inheritance, mutation, and crossover.

#### 2.1 IBM Parameter Initialization and evaluation

We initialized IBM parameters using randomly generated values for 5000 solutions (individuals) based on theoretical viable bounds as follows:

Table 2.1: Randomized initialization ranges for Individual parameters

| **Symbol** | **Bounds** | **Rationale** |
|---|---|---|
| $\kappa$ | $[0.01, 0.99]$ | Parameter value range |
| $v$ | $[0.01, 0.99]$ | Parameter value range |
| $m$ | $[0.01, 0.99]$ if migratory else 0 | Lower limit restricts maximum travel in one time step to $< 90$ distance units at maximum length, upper limit restricts maximum cost to one distance unit to $< 1.$ at maximum length. |
| $l_b$ | $[\frac{\dot{v}}{3}, 0.99]$ | Lower limit restricted to $dl$ at time 1 given unlimited reserve. The smallest possible $l_b$ given the fastest possible daily development. |
| $g$ | $\left[\frac{l_b}{\kappa}, \frac{1}{\kappa}\right]$ | Restricted so that cost to grow 1 unit of volume cannot exceed maximum reserve energy in 1 unit of volume and is greater than the minimum reserve energy density required to achieve birth (if lower than it would not be able to grow at birth). $\left(\frac{[E_G]}{[E_M]} \leq \frac{[E_M]}{[E_M]}\right) = g\kappa \leq 1$ $\left(\frac{[E_G]}{[E_M]} \leq \frac{[E_B]}{[E_M]}\right) = g\kappa \leq l_b$ |

| Symbol | Bounds | Rationale |
|---|---|---|
| $u_H^p$ | $[(1-\kappa)l_b^3, (1-\kappa)]$ | Limits derived from assumptions that maturation thresholds occur at a fixed size $\dot{k}_J = \dot{k}_M$ and $l_b < l_p < 1$ |
| $d_{NB}$ | $[1, 361]$ if migratory else 364 | Upper limit bound by fastest possible migration of 1 day each way, 1 day to reach adulthood, and 1 day to breed. |
| $d_M$ | $\left[1, \frac{365-d_{NB}}{2}\right]$ if migratory else 0 | Upper limit bound by longest possible one way migration given $d_{NB}$ |

Derived parameter values were calculated as follows:

Table 2.2: Derived individual parameters

| Symbol | Value | Rationale |
|---|---|---|
| $u_H^b$ | $(1-\kappa)l_b^3$ | Derived from assumption that maturation thresholds occur at a fixed size $\dot{k}_J = \dot{k}_M$ |
| $\dot{k}_M$ | $\frac{v}{g}$ | Calculated to assume $L_M = 1$ to define limits for g and calculate $a_b$. |
| $a_b$ | $\frac{3}{\dot{k}_M} \int_0^{x_b} \frac{d_x}{(1-x)x^{2/3}(\alpha_b - B_{xb}(\frac{4}{3},0))}$ | Calculated based on DEB embryo development dynamics given $l_b$, $\dot{k}_M$, and minimum possible $e_b$ ($e_b = l_b$) to estimate incubation upper limit to record $u_R$ and $e_b$ at time of egg production $(365 - a_b)$. Re-adjusted at end of IBM for realized $e_b$ for subsequent iterations, although difference in values is generally negligable on the integer scale. See Eq 2.38 in [3] for detail. |
| $u_E^0$ | $\left(\frac{3g}{\alpha_b - B_{xb}(\frac{4}{3},0)}^3\right)$ | Equal to scaled energy cost of one egg. Back calculated from $l_b$ and $e_b$ based on DEB embryo development dynamics. See Eq 2.42 in [3] for detail. |
| $fecundity$ | $\frac{u_{R(365-a_b)}}{u_E^0}$ | |

> **Note**
>
> After the first IBM iteration Initial $e$ is set to the realized $e_b$ from the previous iteration to relax the initial assumption that the parent achieved energetic equilibrium with the environment at time of egg production.

We then evaluated viability and fecundity (fitness) of the solutions with the IBM.

#### 2.2 Optimization procedure

The optimization routine is governed by the following parameters:

Table 2.3: Optimization parameter values were chosen based on simulation tests (see Optimization validation and tuning).

| Parameter | Value | Description |
|---|---|---|
| popsize | 5000 | Number of solutions (individuals) in each iteration |
| generations | 200 | Number of optimization iterations per run |
| replacerate | 1 | Proportion of solutions to replace with selected solutions. |
| crossoverrate | 0.5 | Proportion of new solutions resultant from recombination. |
| mutationrate | 0.5 | Proportion of new solutions resultant from mutation. |

##### 2.2.1 Selection

We only retained solutions for iterative optimization that were viable (alive at the end of IBM), and produced more that 1 offspring ($fecundity \leq 1$). From the viable set of solutions, we selected a new population of 5000 solutions using stochastic universal sampling. This sampling technique selects solutions proportional to their resultant fecundity by using a single random value to sample solutions which reduces biased representation of lower fitness solutions due to stochastic noise [2].

##### 2.2.2 Recombination/Crossover

We then replaced a random half of the solutions with new solutions generated from recombination. For recombination, we pair-wise combined values between solutions by summing a randomized proportion of the value from each parent solution for every parameter value, creating two new solutions from each pair. This mechanism serves to exploit fitness peaks in the solution space found by solutions and combining them to create potentially better-performing solutions. We used probabilistic selection to select pairs based on pair-wise holistic parameter similarity. This technique, niching, prevents dampening exploitation of fit solutions by limiting combination between solutions which have identified disparate peaks in fecundity across a multi-modal fitness landscape.

##### 2.2.3 Mutation

We then replaced a random half of solutions with new solutions generated from mutation. For mutation, we summed parameter values from the solution selected to be mutated with a random proportion of a randomly generated parameter value. This mechanism serves to generate novel solutions for expanded exploration of the sample space in future iterations.

##### 2.2.4 Re-evaluation

We re-evaluated the new set of solutions in the IBM for fecundity and optimized for a total of 200 iterations (generations). To maximize broad exploration of the parameter space we implemented a nested routine such that this process was repeated 1100 times (1100 runs with 200 generations each). Runs were run in parallel to maximize computational efficiency using the 'Futures' package in R.

#### 2.3 Optimization validation and tuning

We used graphical analysis of simulation tests to determine optimization parameter values and validate the optimization algorithm.

We simulated every combination of 3 different values of popsize, replacerate, crossoverrate, and mutation rate for 500 generations for a movement scenario and a non-movement scenario. We ran 5 optimization runs for each combination.

| popsize | replacerate | crossoverrate | mutationrate |
|---|---|---|---|
| 2500 | 0.5 | 0.25 | 0.1 |
| 5000 | 0.75 | 0.5 | 0.25 |
| 10000 | 1 | 0.75 | 0.5 |

We then used a custom shiny app to visualize fecundity max, mean and variance over generation time for each set of runs. In most runs, optimization slowed considerably after ~200 generations but variance between runs prompted us to take a nested approach to optimization, to combat the algorithm getting stuck on local maxima or insufficiently exploring the sample space. This app is available in the code repository on github.

We also visualized parameter values of the top performing solution and the number of unique solutions over generation time. We used visualizations of the parameter values to ensure the routine was exploring the sample space by examining the range of values over generation time and used the visualization of unique solutions overtime to validate crossover and mutation were generating variation in solutions and premature convergence did not occur.

We performed 10 runs in parallel in sets of 100 and compared the top 5 solutions across conditions. To ensure reproducibility, we set seeds for random number generation. An arbitrary seed value was chosen for each set to generate seeds for each run such that the seed for a run was the same for each migratory strategy but different across runs. Graphical comparison of solutions after 300 runs demonstrated that most runs converge on similar solutions (Figure 2.1). We conservatively chose to run an additional 700 runs, which demonstrated a similar visual result (Figure 2.2).

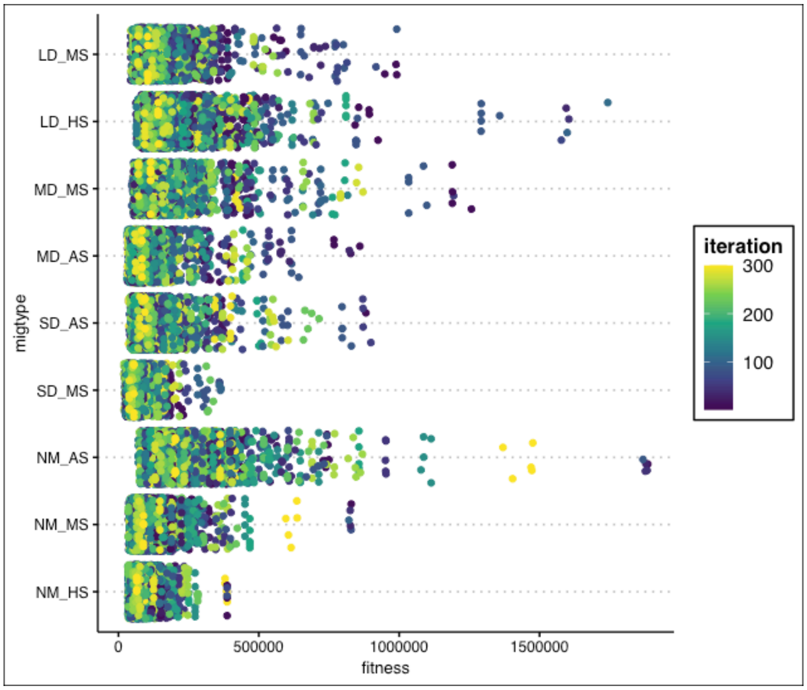

Figure 2.1: Fecundity of solutions per strategy from 300 runs (3 sets of 100)

To further validate relative differences in fecundity across movement scenarios were not an artifact of non-convergence or unequal exploration of the solution space, we performed an additional set of 180 runs per migratory strategy pre-initialized with parameters from five of the highest fecundity solutions for each strategy from the initial 1000 runs to equalize exploration of the solution space between strategies. Pre-initialized runs identified fecundity optimums in the range of randomly initialized runs suggesting similar exploration of the solution space with some improvement on absolute value of optimized fecundity but similar patterns of relative differences between strategies. Figure 2.3.

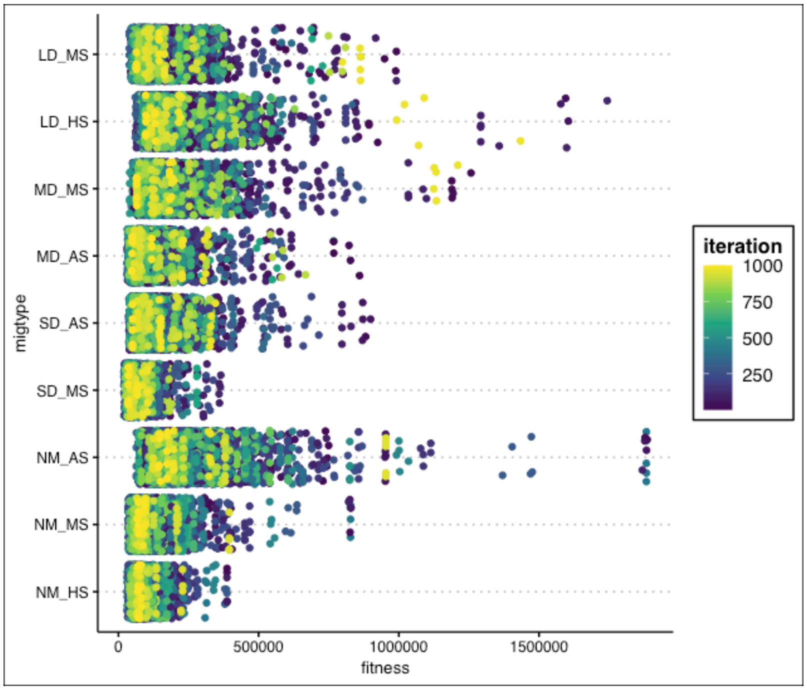

Figure 2.2: Fecundity of solutions per strategy from 1000 runs (10 sets of 100)

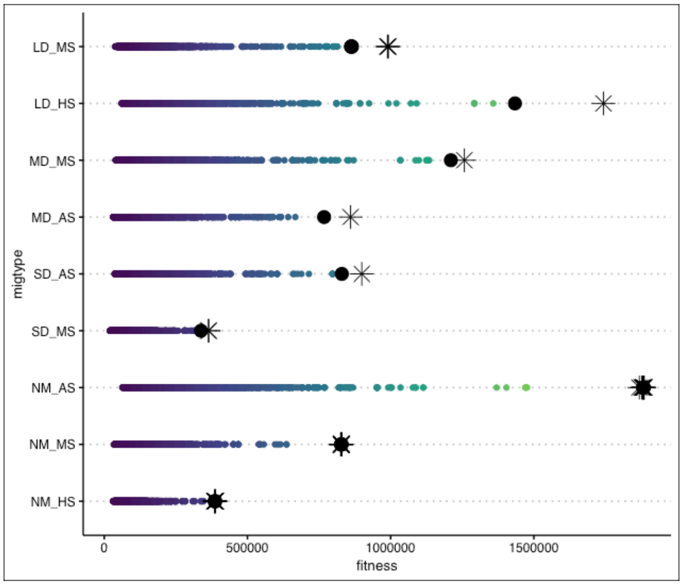

Figure 2.3: Fecundity values of top 1% of solutions from intial 1000 runs shown as large black dots and pre-initialized runs shown aslarge black asterisks. Overlapping colored dots show fecundity values for all solutions from initial 1000 runs.

### Bibliography

[1] Volker Grimm et al. "The ODD Protocol for Describing Agent-Based and Other Simulation Models: A Second Update to Improve Clarity, Replication, and Structural Realism". In: *Journal of Artificial Societies and Social Simulation* 23.2 (2020). DOI: 10.18564/jasss.4259. URL: http://dx.doi.org/10.18564/jasss.4259.

[2] *Introduction to Genetic Algorithms*. Springer Berlin Heidelberg, 2008. DOI: 10.1007/978-3-540-73190-0. URL: http://dx.doi.org/10.1007/978-3-540-73190-0.

[3] Bas Kooijman. "Dynamic Energy Budget Theory for Metabolic Organisation". In: (Oct. 15, 2009). DOI: 10.1017/cbo9780511805400. URL: http://dx.doi.org/10.1017/cbo9780511805400.
