## Appendix S2 (Supplemental Figures) for "Seasonal migration alters energetic trade-off optimization and shapes life history" (from another version of this preprint: version 1, 2023-09-15)

### Appendix S2. Supplementary analysis figures

Allison K. Pierce, Scott W. Yanco, & Michael B. Wunder. Seasonal migration alters energetic trade-off optimization and shapes life history.

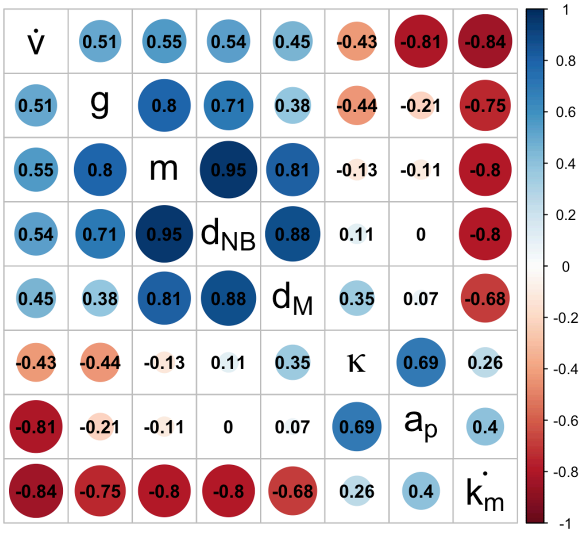

**Figure S1.** IBM parameter correlation matrix.

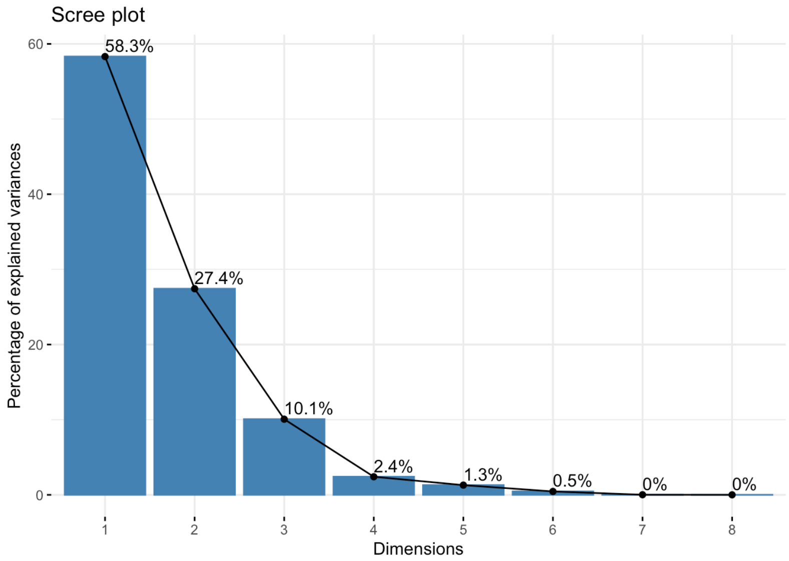


**Figure S2.** PCA Scree plot demonstrating the first 3 axes of variation explained 89.9% of the variance in the optimized IBM parameter data.

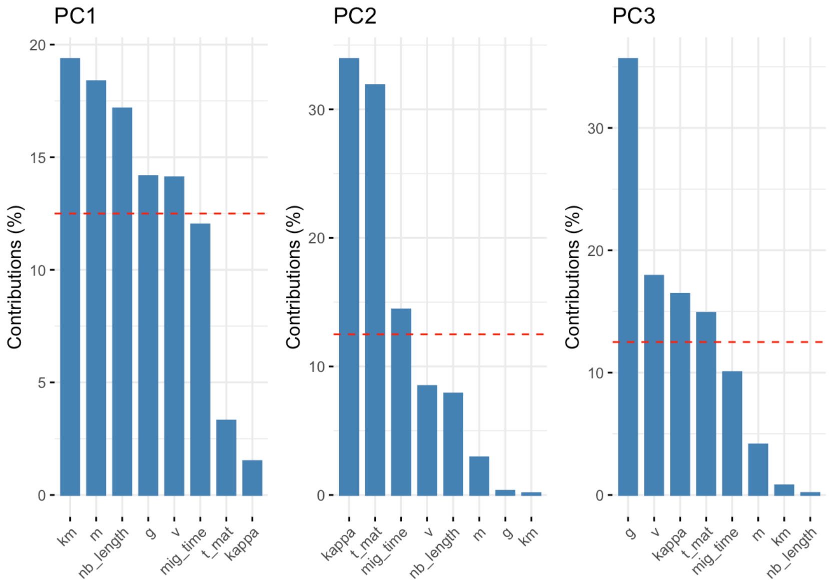


**Figure S3.** Relative contributions of each parameter for PC1, PC2, and PC3 (left to right). Dashed lines indicate the expected average contribution.

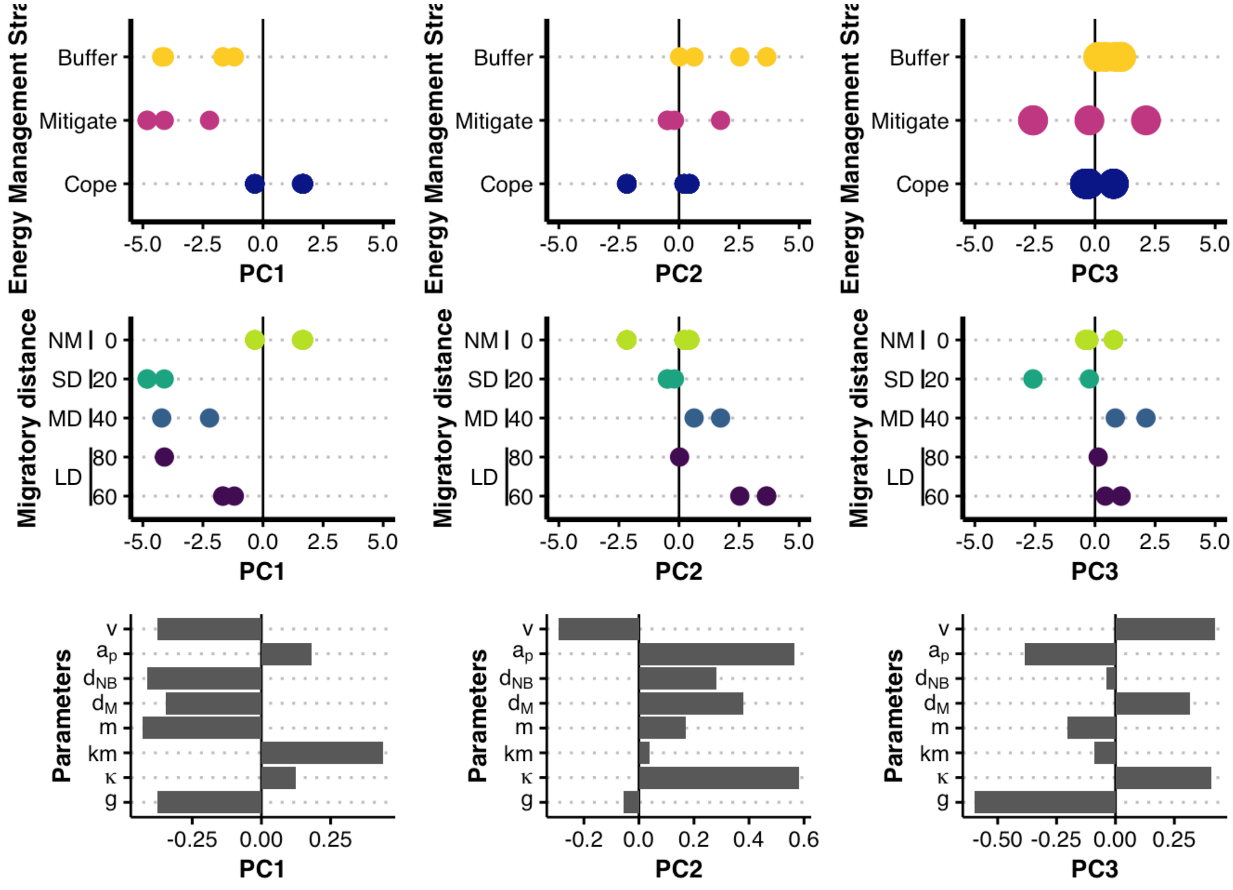


**Figure S4.** Individual PCA data for PC1, PC2, and PC3 plotted by energy management strategy and migratory distance(top) with parameter loadings (bottom).

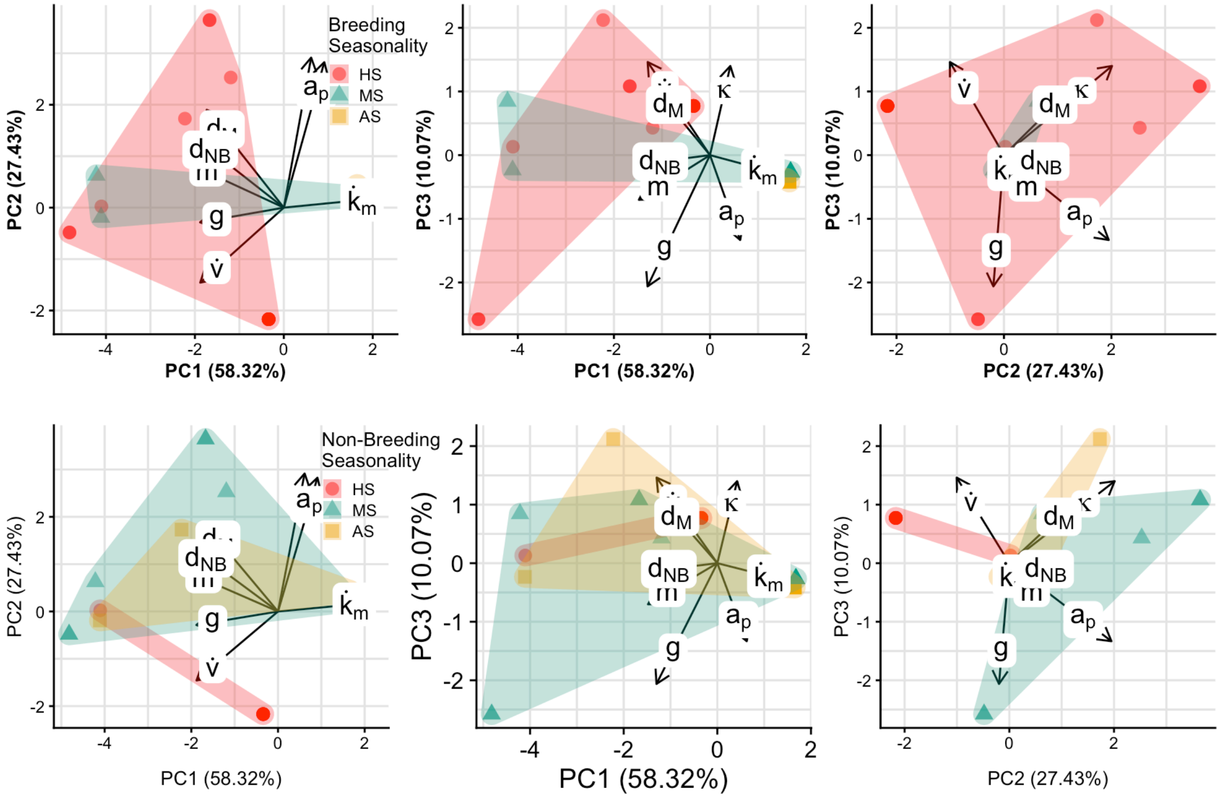


**Figure S5.** PCA Bi-plots colored by breeding (top) and non-breeding seasonality (bottom).

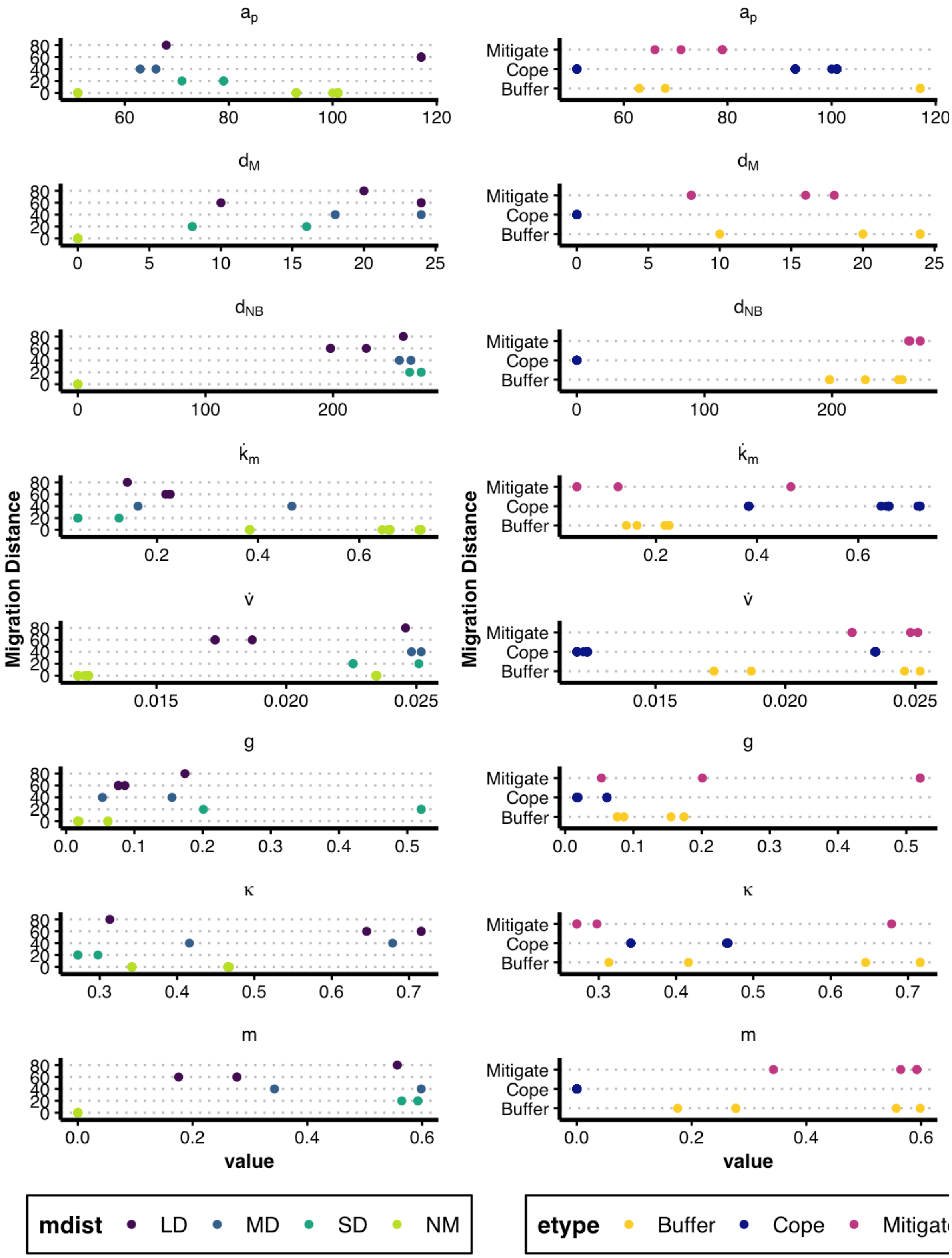


**Figure S6.** Parameter values by migration distance and energy management strategy.

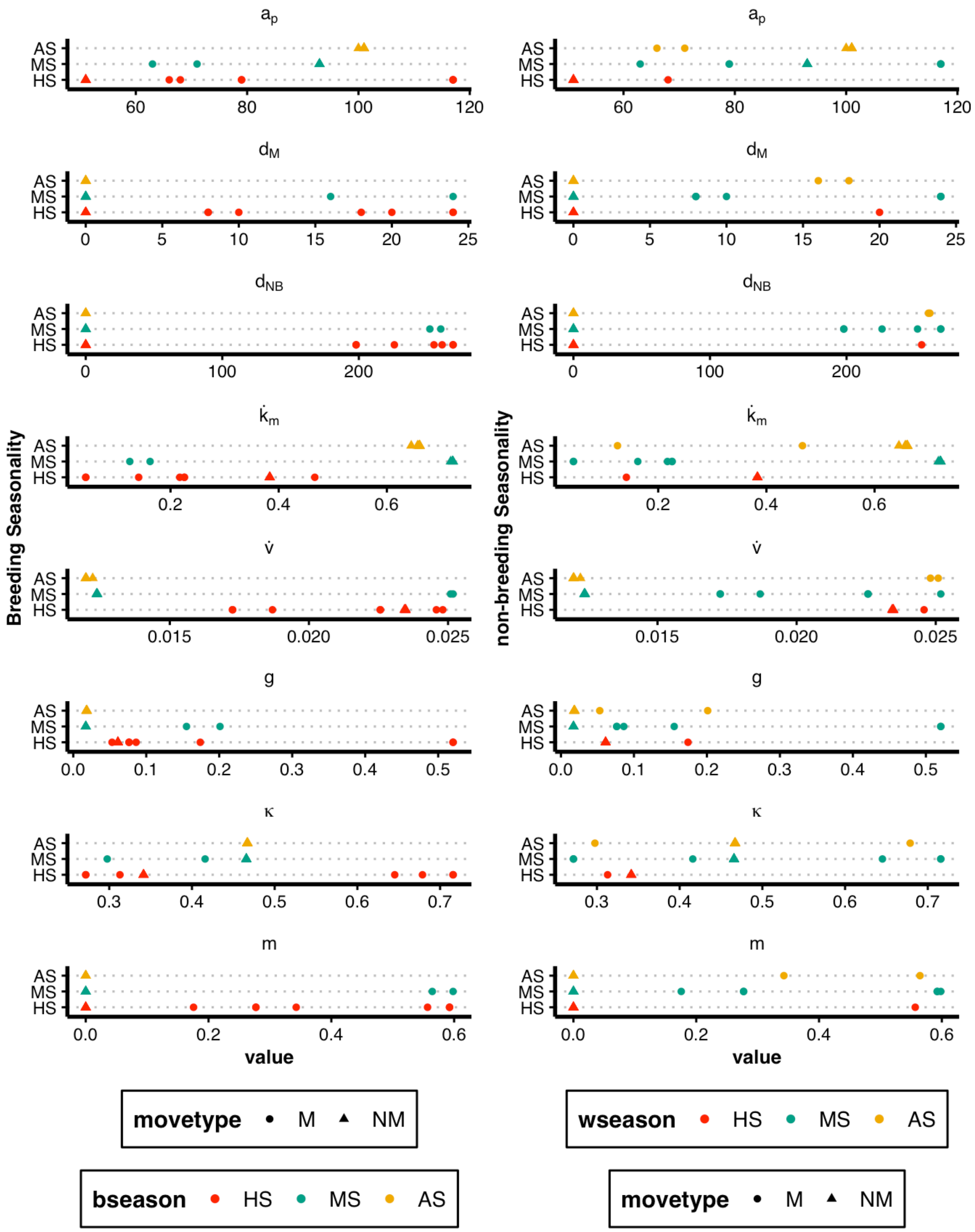


**Figure S7.** Parameter values by breeding and non-breeding seasonality.
